# Pangenome analysis of the soil-borne fungal phytopathogen *Rhizoctonia solani* and development of a comprehensive web resource: RsolaniDB

**DOI:** 10.1101/2020.12.18.423518

**Authors:** A. Kaushik, D.P. Roberts, A. Ramaprasad, S. Mfarrej, Mridul Nair, D.K. Lakshman, A. Pain

## Abstract

*Rhizoctonia solani* is a collective group of genetically and pathologically diverse basidiomycetous fungus that damages economically important crops. Its isolates are classified into 13 Anastomosis Groups (AGs) and subgroups having distinctive morphology and host range. The genetic factors driving the unique features of *R. solani* pathology are not well characterized due to the limited availability of its annotated genomes. Therefore, we performed genome sequencing, assembly, annotation and functional analysis of 12 *R. solani* isolates covering 7 AGs and selected subgroups (AG1-IA, AG1-IB, AG1-IC, AG2-2IIIB, AG3-PT (isolates Rhs 1AP and the hypovirulent Rhs1A1), AG3-TB, AG4-HG-I (isolates Rs23 and R118-11), AG5, AG6, and AG8), in which six genomes are reported for the first time, wherein we discovered unique and shared secretomes, CAZymes, and effectors across the AGs. Using a pangenome comparative analysis of 12 *R. solani* isolates and 15 other basidiomycetes, we also elucidated the molecular factors potentially involved in determining the AG-specific host preference, and the attributes distinguishing them from other Basidiomycetes. Finally, we present the largest repertoire of *R. solani* genomes and their annotated components as a comprehensive database, viz. RsolaniDB, with tools for large-scale data mining, functional enrichment and sequence analysis not available with other state-of-the-art platforms, to assist mycologists in formulating new hypotheses.

## Introduction

*Rhizoctonia solani* Kühn (teleomorph: *Thanatephorus cucumeris* [Frank] Donk) is considered as one of the most destructive soil borne plant pathogens causing various diseases including pre- and post-emergence damping-off of seedlings, crown and root rots, black scurf of potato, take-all of wheat, sheath blight of rice and maize, brown patch of turf, and postharvest fruit rots (1, 2). This necrotrophic fungus infects a wide range of economically important plant species, belonging to more than 32 plant families and 188 genera, and is responsible for 15% to 50% agricultural damages annually (3). Broadly, it is classified among 13 Anastomosis Groups (AGs) with distinctive morphology, physiology, pathogenicity host range, and highly divergent genetic composition (4). Most *R. solani* AGs are further divided into subgroups, also called IntraSpecific Groups (ISGs), which differ in pathogenicity, virulence, ability to form sclerotia, growth rate, and host range preference (5). Although field isolates of *Rhizoctonia* infected plants are usually found to be infested with one or more AGs, each AG subgroup can still have its own host preference. For instance, *Arabidopsis thaliana,* was found to be susceptible to AG2-1 sub-group isolates but resistant to AG8 isolates (6), which suggests that genetic divergence is the inherent characteristic of *Rhizoctonia* species.

Over the last two decades, our understanding of the genetic divergence among different *R. solani* AGs has improved to the point that it is now evident that all AGs and their sub-groups are genetically isolated, non-interbreeding populations (7). The rapid and relatively low-cost of generation of genomic sequences and other *‘omics’* datasets has played a significant role in furthering our understanding of the host-pathogen interactions and ecology of *Rhizoctonia* species. (8–12). The analysis of these genomic sequences and functional components revealed several novel or previously unrecognized classes of *R. solani* genes among different AGs that are involved in pathogenesis in a host-specific manner, e.g. effector proteins and carbohydrate-active enzymes (CAZymes) (13). Additionally, analysis of differentially expressed genes in different isolates has enabled researchers to predict the adaptive behavior of this fungus in different hosts and the associated virulence (14, 15). However, the majority of this information has come from the analysis of isolates belonging to only a small number of AGs for which complete genome and/or transcriptome sequences are available. In fact, until now, draft genome assemblies belonging to only 4 of the 13 AGs have been reported viz. AG1-IA (16), AG1-IB (17), AG2-2IIIB (13), AG3-Rhs1AP (18), AG3-PT isolate Ben-3 (19) and AG8 (20). This limited availability of genome sequences and the predicted proteomes across the 13 different AGs and their subgroups is one of the important barriers hindering the understanding of functional complexity and temporal dynamics in *R. solani* AGs and their subgroups.

In this study, we report whole-genome sequencing, assembly and annotation of 12 *Rhizoctonia* isolates from 7 AGs; of which genome sequences of three AGs (AG4, AG5, and AG6), two subgroups (AG1-IC and AG3-TB {or AG3-T5}) and a hypovirulent isolate (AG3-1A1) of the subgroup AG3-PT are being reported for the first time. The draft genome of the AG3-PT isolate 1AP (alternatively named as Rhs1AP) was previously reported (Cubeta et. al., 2014) (18), but was re-sequenced for comparative purposes, as AG3-1AP. Furthermore, to understand genetic diversity among different *R. solani* isolates, we performed inter-proteome comparative analyses, including ortholog analysis at the pangenome level and protein domains profiling for secreted components, virulent proteins, and CAZymes in all 12 *R. solani* isolates. To make these high-quality draft *R. solani* genomes and features readily accessible to a broad audience of researchers, we built a comprehensive and dedicated web resource, viz. RsolaniDB, for hosting and analyzing the available genomic information predicted at the transcript-, and protein-level in different *R. solani* AGs. The presented web-resource includes detailed information on each *R. solani* isolate, such as the genome properties, predicted gene, transcript and protein sequences, predicted gene function, and protein orthologues among other AG sub-groups, along with tools for Gene Ontology (GO) and pathway enrichment analysis, sequence analysis, and visualization of gene models.

## Materials and Methods

### Isolation of genomic DNAs for sequencing

Details regarding *R. solani* isolates used for sequence analyses are presented in Table S1 and S2. Fungal cultures were purified by the hyphal tip excision method (21) and maintained by sub-culturing on potato dextrose agar (PDA, Sigma Aldrich catalog # P2182, St. Louis, MO, USA). The PDA was amended with kanamycin (25 μg/ml) and streptomycin (50 μg/ml) to inhibit bacterial growth. Isolates were grown in Potato Dextrose Broth (PDB, Sigma Aldrich catalog # P6685) broth at 100 rpm and 25 C for 4 to 6 days, mycelia collected by filtration through 2 layers of sterile cheese cloth, washed 2 X with sterile distilled water, gently squeezed and placed on 4 layers of paper towel to remove surface water, and then snap-frozen in liquid nitrogen and stored at −80 C till use. Genomic DNA was extracted from mycelia using both the CTAB method (22) and a protocol recommended by the manufacturer (User-Developed Protocol: Isolation of genomic DNA from plants and filamentous fungi using the QIAGEN® Genomic-tip, Qiagen Inc.). RNA was extracted from fungal isolates and from tobacco detached leaves infected with corresponding fungal isolates, using the Qiagen RNeasy Plant Mini Kit (Qiagen Inc. Germantown, MD, USA). Extracted genomic DNA and RNA was quantified with a Qubit Flex Fluorometer (Thermo Fisher Scientific, Waltham, MA, USA). AG and subgroup identity of the fungal isolates was verified by ITS-PCR, sequencing and homology analysis with nucleotide sequences available in the NCBI database (23).

### RNA extraction

*Nicotina tabacum* seedlings were raised to four-leaf stage on potting mix (Pro-mix, Premier Horticulture, USA) in the greenhouse at ambient temperature (22° - 24° C) and with four hours supplemental light with a mercury lamp. Two leaves were excised from each seedling and placed on a tray on two piece of wet paper towels. For inoculation, seven to eight agar plugs from the margin of fresh *R. solani* growth on 1/4^th^ concentration of PDA (potato dextrose agar) were placed on the adaxial surface of each leaf. For control, only seven to eight agar plugs from 1/4^th^ PDA were placed. Each tray was closed with a lid and incubated on lab bench at ambient temperature and light.

After 5 days, yellow to necrotic symptoms were noticeable on *R. solani* treated leaves but no symptoms appeared on control leaves surrounding the plugs. The control and infected patches were excised with a sterile scalpel, snap frozen in liquid nitrogen and processed for RNA extraction with RNeasy Plus Mini Kit in RLC buffer (Qiagen Sciences Inc., Germantown, MD, USA). The purified RNA was treated with DNase at 37° C for 30 min, extracted with phenol and Phenol: chloroform, precipitated with ethanol, and dissolved in RNase-free water.

### Construction of genomic and RNA libraries and sequencing

For making genomic libraries, an input of 500ng of DNA from each sample was sheared on Covaris (Covaries E series) and paired-end libraries were prepared for sequencing using Illumina’s HiSeq 2000 platform. From end repair until adapter ligation and purification steps of the paired-end libraries were prepared using the protocol “Illumina library prep” on the IP-Star automated platform from Diagenonde (Diagenode IP Star) as per the manufacturer’s protocol.

Post ligation, manual protocols were used for gel size selection and PCR amplification using the standard Illumina PCR Cycle (Kapa high-fidelity master mix). The prepared libraries were analyzed on bioanalyzer and quantified using Qubit (Thermo Fisher). The normalized libraries were pooled for sequencing (insert size of 500bp) and submitted for HiSeq 2000 sequencing at Bioscience Core Laboratory of King Abdullah University of Science and Technology. Strand-specific mRNA sequencing was performed from total RNA using TruSeq Stranded mRNA Sample Prep Kit LT (Illumina) according to manufacturer’s instructions. Briefly, polyA+ mRNA was purified from total RNA using oligo-dT dynabead selection. First strand cDNA was synthesised using randomly primed oligos followed by second strand synthesis where dUTPs were incorporated to achieve strand-specificity. The cDNA was adapter-ligated and the libraries amplified by PCR. Libraries were sequenced in Illumina Hiseq2000 with paired-end 100bp read chemistry.

### *De novo* assembly, genome annotation and bioinformatic analysis

#### Data preprocessing

Adapter sequences in genomic reads in FASTQ format were trimmed using the trimmomatic tool (version 0.35) (24), followed by trimming low-quality bases at read ends. Read quality was evaluated using the fastqc tool (version 0.11.8) (25). Reads with length < 20 bp and average quality score < 30 were also removed. For genome heterogeneity analysis, *k-mer* distribution analysis on resulting DNAseq reads was performed using jellyfish (version 2.2.10) (26), which estimated best *k-mer* length for each genome. Histogram distributions of different *k-mers* for the best *k-mer* length was plotted using the *-histo* module of the jellyfish program. In addition, the available raw RNAseq paired-end reads (Table S2) were quality trimmed and preprocessed using with the same approach used for DNAseq reads. The quality trimmed reads were then subjected to *denovo* assembly using Trinity which predicted transcript sequences (27).

#### Genome assembly

Quality trimmed reads were subjected to *denovo* genome assembly using SPAdes (version 3.7.0) in which a defined range of *k-mer* lengths (21,33,55,65,77,101 and 111) was used for contig formation (28). Quast (version 4.5) was used for quality evaluation of predicted contigs (29). Scaffolds were subsequently predicted from contigs using SSPACE (version3.0) (30) and gaps in assembled scaffolds filled using five consecutive runs of GapCloser (version 1.12) (31). For samples with RNAseq dataset available, genome scaffolding was further improved using the Rascaf program (32). Genome quality was evaluated with BUSCO (version 3.0.1) (33) and scaffolds subjected to ITSx (version 1.1) (34) for ITS sequence prediction. Thereafter, phylogenetic tree was constructed with megax software (35) using the neighborhood joining method (10000 bootstraps), in which ITS2 sequences were aligned using ClustalW (36). The resulting tree was saved in the newick format and visualized together using Phylogeny.IO (37) and ETE toolkit (38). Redundans python script was then used to predict the homozygous genome by reducing the unwanted redundancy to improve draft genome quality (39). Resulting scaffolds were aligned with mitochondrial genomes of *R. solani* and other Basidiomycota using blastn program (version 2.6.0; e-value ≤ 1e^-5^) (40) and mapped mitochondrial contigs were removed to retain only the nuclear genome for subsequent annotation.

#### Genome annotation

The draft genome was annotated using the MAKER (version 2.31.8) pipeline(41), which predicted intron/exon boundaries, transcript and protein sequences. For the annotation, repeat regions were masked using RepeatMasker (version 4.0.5; model_org=fungi) (42). Protein homology evidence was taken from UniProt protein sequences (Reviewed; family: Basidiomycota) (43). For EST evidences, RNAseq reads were assembled into transcripts using Trinity *denovo* assembler (version 2.0.6) (27). For genomic datasets without corresponding RNAseq datasets available, the EST sequences of alternate organisms were used from previously published *R. solani* genome annotations viz. AG1-IA (16), AG1-IB (17), AG2-2IIIB (13), AG3-Rhs1AP (18), AG3-PT isolate Ben-3 (19) and AG8 (20). The functional domains, PANTHER pathways (44) and Gene Ontology (GO) terms (45) in the predicted protein sequences were assigned using InterProScan (version 5.45-80.0) standalone program (46). The functional domains assigned to each protein included the information from ProSiteProfiles (47), CDD (48), Pfam (49) and TIGRFAMs (50), resulting in the annotated genome in GFF3 format using iprscan2gff3 and ipr_update_gff programs (46).

The fungal AROM protein sequences were identified by mapping the *R. solani* proteome on pentafunctional AROM polypeptide sequences from UniProt (organism Fungi) using blastp (e-value ≤ 0.001) (40). The resulting candidate AROM sequences in each *R. solani* proteome were analyzed using HMMER webserver (51). We also identified the predicted secreted proteins in each of the *R. solani* proteomes using signalp (version 5.0) (52). For identification of proteins with a transmembrane domain phobius (version 1.01) (53) was used. We used targetp (version 1.1) to predict proteins with mitochondrial signal peptides (54). However, since we already removed mitochondrial contigs from assembled genomes, we did not observe any proteins with a mitochondrial signal peptide. Effector proteins in each *R. solani* secretome were predicted using effectorP webserver (version 2.0) (55). The Carbohydrate Active enZyme (CAZyme) in *R. solani* proteomes were predicted using dbCAN2 webserver, in which only the proteins predicted by at least two prediction methods were considered (56). The CAZyme family predicted by HMMER was used for the selected proteins.

#### Orthology

Orthologous proteins across all proteomes were identified with orthoMCL clustering using the Synima program (57, 58), which identified core, unique and auxiliary regions in each *R. solani* proteome. This program was also used for predicting genome synteny using inter-proteome sequence similarity. ShinyCircos was used for circular visualization of synteny plots (59, 60).

### RsolaniDB database development

The RsolaniDB (RDB) database was built to host *R. solani* reference genomes, transcript and protein sequences in FASTA format, along with genome annotations included in GFF3 format. For each genome, the information in the database was structured as entries, in which each entry included a list of details about a given transcript and protein, i.e., intron-exon boundaries; predicted functions; associated pathways and GO terms; predicted sequences; orthologs and functional protein sequence domains predicted from InterPro, PrositeProfile and Pfam. The identifier format for each entry (i.e., RDB ID) start with ‘RS_’ and AG subgroup name followed by a unique number. We also included five previously published *R. solani* annotated genome sequences (i.e., AG1-1A, AG1-1B, AG2-2IIIB, AG3-PT and AG8) with their gene identifiers converted into the RDB ID format. The database was written using DHTML and CGI-BIN Perl and MySQL language, to allow users perform list of tasks, including text-based search for the entire database; or in AG-specific manner. We also included a list of tools to assist users in performing number of down-stream analysis, including RDB ID to protein/transcript sequence conversion; FASTA sequence-based BLAST search on entire database or AG-specific manner; tool to retrieve orthologs for a given set of RDB IDs along with tools for functional enrichment analysis. The GO-based functional enrichment tool for gene set analysis of given RDB IDs was build using topGO R package (61). Whereas the pathway-based gene set analysis was developed to predict significantly enriched PANTHER pathway IDs for a given set of RDB IDs.

## Results

### Genome-wide comparative analysis of *R. solani* assemblies and its annotation

We performed the high-depth sequencing, *denovo* genome assembly and annotation of 12 *R. solani* isolates. For qualitative evaluation of these assemblies, we used genome sequences of a basidiomycetous mycorrhizal fungus *Tulasnella calospora* (Joint Genome Institute fungal genome portal MycoCosm (http://genome.jgi.doe.gov/Tulca1/Tulca1.home.htm) and *R. solani* AG3-PT as negative and positive controls, respectively. Overall, the draft genome assemblies of the *R. solani* isolates shows remarkable differences in the genome size, ranging anywhere from the smaller AG1-IC (~33 Mbp) to the larger AG3-1A1 (~71 Mbp) isolate genomes (Table S3). The number of contigs generated are also highly variable ranging between 678-11,793, in which the newly reported assemblies of AG1-IC and AG3-T5 has highest N50 lengths of 1,00,597 bp and 1,96,000 bp respectively (Table S3). The heterogeneity in genomic reads was predicted by analyzing the distribution of different *k-mers* in *R. solani* genomic sequencing reads. The analysis reveals a shoulder peak along with the major peak in *k*-mer frequencies for AG2-2IIIB, AG3-1A1, AG3-1AP and AG8, indicating the possible heterogeneity of these genomic reads of these isolates (Figure S1). The G+C content ranged from 47.47% to 49.07%, with a mean of 48.43% (Table S3). The quality of these draft genomes was evaluated using BUSCO with scores ranging between ~88-96% (Table S3), indicating the completeness of essential fungal genes in the predicted assemblies. In order to evaluate the reliability of the genome assemblies, we further compared our draft genomes with previously published assemblies of *R. solani* isolates, i.e., AG1-IA, AG1-IB, AG2-2IIIB, and AG8 (Figure S2). The mummer plot (62) comparison shows the overall co-linearity and high similarity among similar assemblies, wherein AG8 assemblies are least co-linear, possibly due to the heterokaryotic nature of the AG8 genome (20, 63). Among the presented draft genome sequences, a large number of syntenic relationships (Figure 1A) are also identified (length > 40,000bp), wherein all the given isolates share at least four highly similar syntenic region, except *T. calospora* (outgroup), which does not share any syntenic region with *R. solani* isolated for the given threshold of > 40,000 bp (Figure 1B). Similarly, our analysis shows that AG5, AG2-2IIIB and AG3-1A1 shares comparatively lower syntenic regions, whereas AG3-PT (positive control) shares highest number of syntenic regions with other *R. solani* isolates. In fact, we observed that most of the closely related AGs share large number of syntenic relationships, e.g., high similarity among AG3 sub-groups. Overall, the analysis exhibits the first line of evidence that indicates widespread collinearity and regions of large similarity across genetically distinct isolates, with *T. calospora* as an outlier.

**Figure 1.**
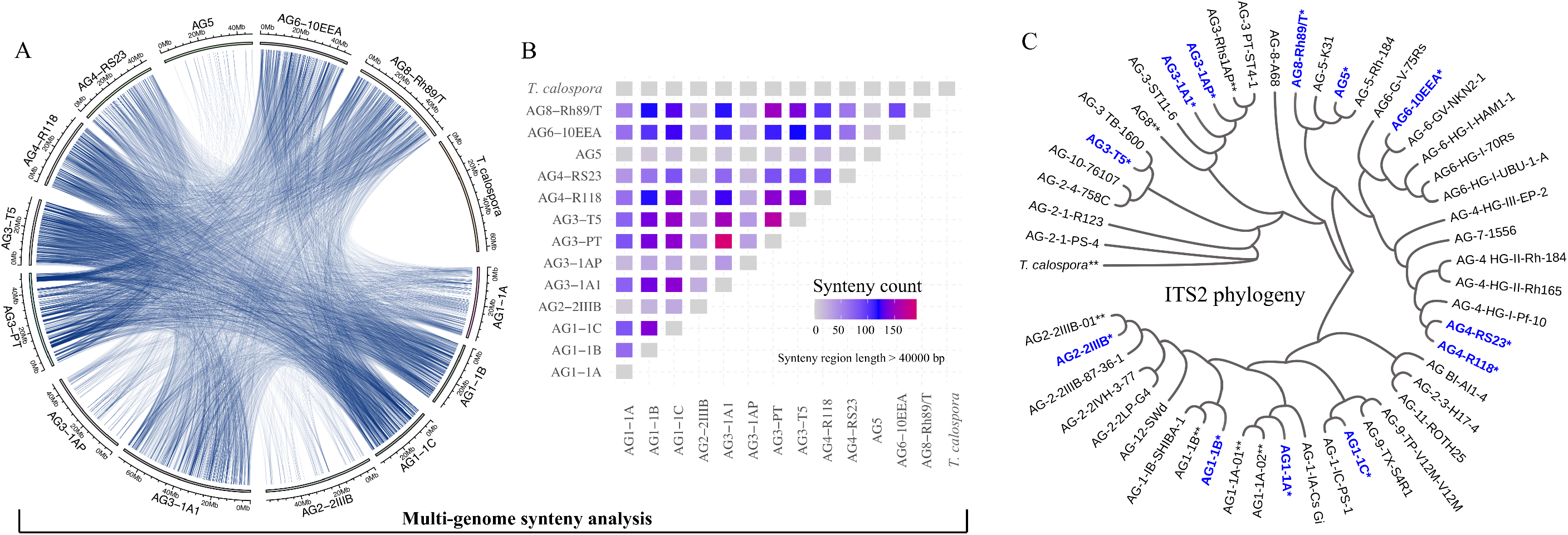
A. **Circos plot**. The Circos plot represents the syntenic relationship between genomes of the different AGs of *Rhizoctonia solani* Kühn. Each line represents the region of genomic similarity predicted with Synima. Only the regions with coverage > 40,000 bases were enumerated and shown. B. The plot highlights the number of high-similarity syntenic regions (coverage > 40,000 bp) shared between each pair of genomes, including *T. calospora.* The red connection represents corresponding isolates sharing comparatively large number of syntenic relationships than other pair of isolates. Here, self-hits were removed or not shown. C. **ITS2 phylogeny**. ITS2 sequences of the tester strain were obtained from the NCBI database and were clustered with ITS2 sequences from assembled *R. solani* genomes (highlighted with blue color and *), along with ITS2 sequences from previously published *R. solani* genome assemblies (marked with **). The phylogenetic tree was constructed using megax software with 10,000 bootstrapping steps (see methods), after which resulting tree and corresponding alignment were visualized together using Phylogeny.IO.

Subsequently, we performed the ITS2-based phylogeny to compare the ITS2 sequences of the 12 newly sequenced *R. solani* isolates with that of the known *R. solani* tester strains (as positive controls) and *T. calospora* as an outgroup (Figure 1C), wherein for AG3-PT, we were not able not predict the ITS sequences. The observed phylogenetic clusters of AGs reflect strong similarity in ITS2 sequences of assembled genomes with respect to that of tester strains of *R. solani.* For instance, the AG1-IA cluster includes four strains, all belonging to same AG, *i.e.*, AG1-IA. Similarly, ITS2 sequences of different AG3 and AG4 subgroups are clustered within their respective clade, whereas the outgroup *T. calospora* shows distinct architecture, providing strong evidence in favor of the correct methods used for genome assemblies. Intriguingly, the ITS2 sequences of AG8 subgroup shows remarkable differences, in which sequence of tester strain (i.e., AG-8-A68), previously published genome sequence (i.e., AG8-01) and from the reported genome of this study (i.e., AG8-Rh89/T) are clustered across different clade of the phylogenetic tree.

One of the important proteins known to be strongly associated with fungal evolution and virulence is the penta-functional AROM sequence, with characteristic five domains (64). Here, we characterized the AROM protein sequences in the predicted proteome of all given assemblies (Figure S3). We observed at least one penta-functional AROM sequence in each of the assemblies, in which sequence(s) are present in either complete or partial form. Interestingly, AG3-1A1 has two complete penta-functional AROM protein sequences, a characteristic not observed in any other AG. In AG5, two partial AROM sequences are observed that together completed all five domains observed in the complete penta-functional AROM sequence (65). Although, all assemblies are found to have contiguous AROM protein sequences, the partial AROM sequences in AG5 may represent a fragmented region of the genome assembly and, therefore, warrants further experimental investigations and genome assembly improvements.

### Genome-wide orthologous protein clustering and functional analysis

Intron/exon and transcript boundaries were identified using the maker pipeline (see materials and methods), which predicted 7,394 to 10,958 protein coding transcripts per genome (excluding *T. calospora,* Figure S4) in which AG3-1A1 genome has the highest number of transcripts. Next, using OrthoMCL, the translated protein sequences in all genomes were clustered into the orthologous groups, where each cluster of proteins represented a set of similar sequences likely to represent a protein family. The similarities among the given isolates were enumerated by measuring proteins shared by different proteomes in the same orthoMCL clusters (Figure 2A). As expected, this analysis clearly outgroup *T. calospora*, indicating that it has a different protein family composition than *R. solani* isolates. Although, AG1 and AG4 subgroups, AG3-1A1 and AG3-1AP shows expected similarities and share similar clustering profiles, AG3-PT and AG5 shows a divergent profile of protein families with respect to the other AGs under study. Nevertheless, a large set of orthoMCL clusters share proteins from all/most of the *R. solani* isolates which further indicates inherent similarities as well as unique attributes across these pathologically diverse groups of fungi. For instance, more than 1,400 orthoMCL clusters are composed of proteins belonging to only two AGs, whereas >1,500 clusters are composed of proteins from all 13 *R. solani* isolates and *T. calospora* (Figure 2B). It is expected that these conserved clusters are composed of proteins from core gene families with essential functions, whereas other clusters may host proteins with unique AG-specific roles (Figure 2C). The analysis reveals that AG1-1C, AG2-2IIIB, AG5, AG6-10EEA and AG8 are composed of large number of unique proteins (>1,000 proteins), whereas AG3-1A1 has the highest number of core and auxiliary proteins. The pair-wise comparison of the number of clusters shared by any two AGs highlighted that AG3-1AP shares the highest number of orthoMCL clusters with AG3-1A1, a sector derived hypo-virulent isolate of AG3-1AP (Figure 2D) (66). In fact, AG3-1A1 proteins shares large number of clusters with few other AG subgroups too, including AG1-1C, AG6-10EEA, AG2-2IIIB and AG4-R118.

**Figure 2.**
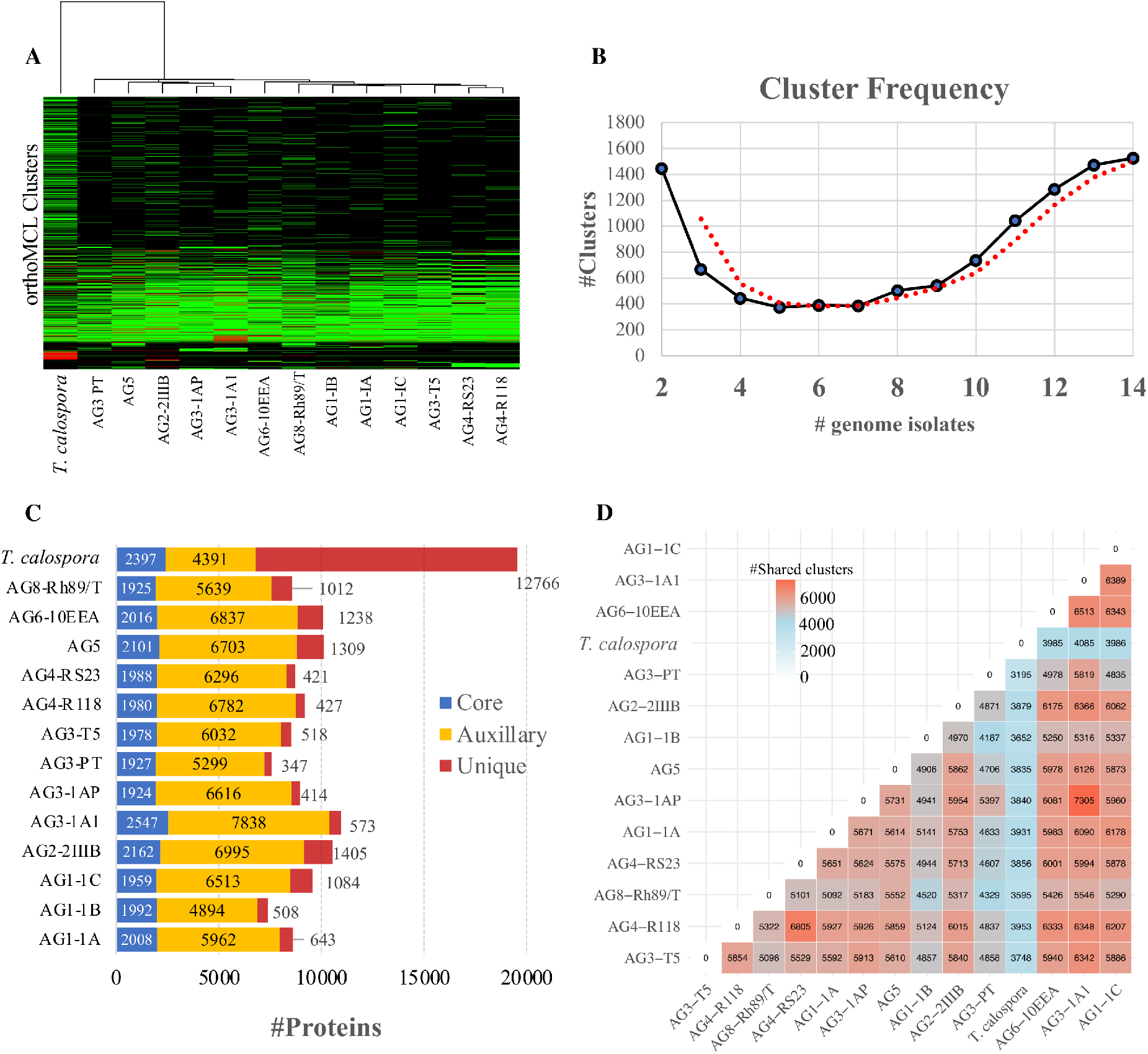
orthoMCL clustering of the predicted proteomes in *R. solani* AGs. A. Heatmap showing protein conservation across all sequenced *R. solani* AGs and *T. calospora*. Each row represents one orthoMCL cluster, and color is proportional to the number of protein members shared within a given cluster from the given species (black: no member protein present; red: large number of protein members present). The hierarchical clustering (hclust; method: complete) analysis enumerates the similarities between different fungal isolates based on proteins shared by them across all orthoMCL clusters. B. **Cluster frequency**. The line plot represents the number of orthoMCL clusters shared by different fungal isolates used in this study. Example, > 1400 orthoMCL clusters are shared by 14 different fungal isolates (including positive and negative controls) used in this study. The bimodal nature of plot represent high similarities across independent proteomes as large number of clusters shares protein members from ≥13 fungal isolates. The red line represents the smoothed curves after averaging out the number of clusters. C. **Protein classification based on the orthoMCL clusters**. The “core” proteins represent the sub-set of proteomes (from each *R. solani* AG and *T. calospsora)* with conserved profile across all the isolates. Similarly, the “unique” sets represent the isolate-specific protein subset. The rest of the protein subsets make the “Auxillary” proteome which are conserved in a limited number of isolates. D. **Shared orthoMCL clusters**. The number of orthoMCL clusters shared between any two isolates. A shared cluster means, a given orthoMCL cluster contains proteins from both the isolates.

To investigate the functional composition of proteins using orthologous groups information, we performed InterPro domain family analysis of proteomes from each AG (Figure S5). Interestingly, the core proteome of most AGs is composed of ~2,000 InterPro domain families, whereas the unique proteome per AG ranged between 101 (for AG3-PT) to 628 (for AG3-1A1). Wherein, the most common protein family that made the unique proteome of *R. solani* subgroups is “Cytochrome P450”, which is essential for fungal adaptations to diverse ecological niches (67) (Figure 3). Similarly, proteins with WD40 repeats are found to be the most common set of the unique proteome in most AGs. In addition, few of the AG subgroups are found to be enriched with a protein family that is significantly associated with its unique proteome only, possibly being involved in the survival of that AG in respective hosts. For instance, the AG1-IB unique proteome is enriched with “NADH: Flavin Oxidoreductase/ NADH oxidase (N-terminal)”, similarly AG3-1A1 is enriched with “ABC transporter-like” and “Aminoacyl-tRNA synthetase (class-II)” InterPro domains. Whereas AG3-PT is found to be uniquely enriched with “Ribosomal protein S4/S9” and AG3-1A1 is uniquely enriched with “Multicopper oxidase (Type 2)” and “Patatin like phospholipase domain”.

**Figure 3.**
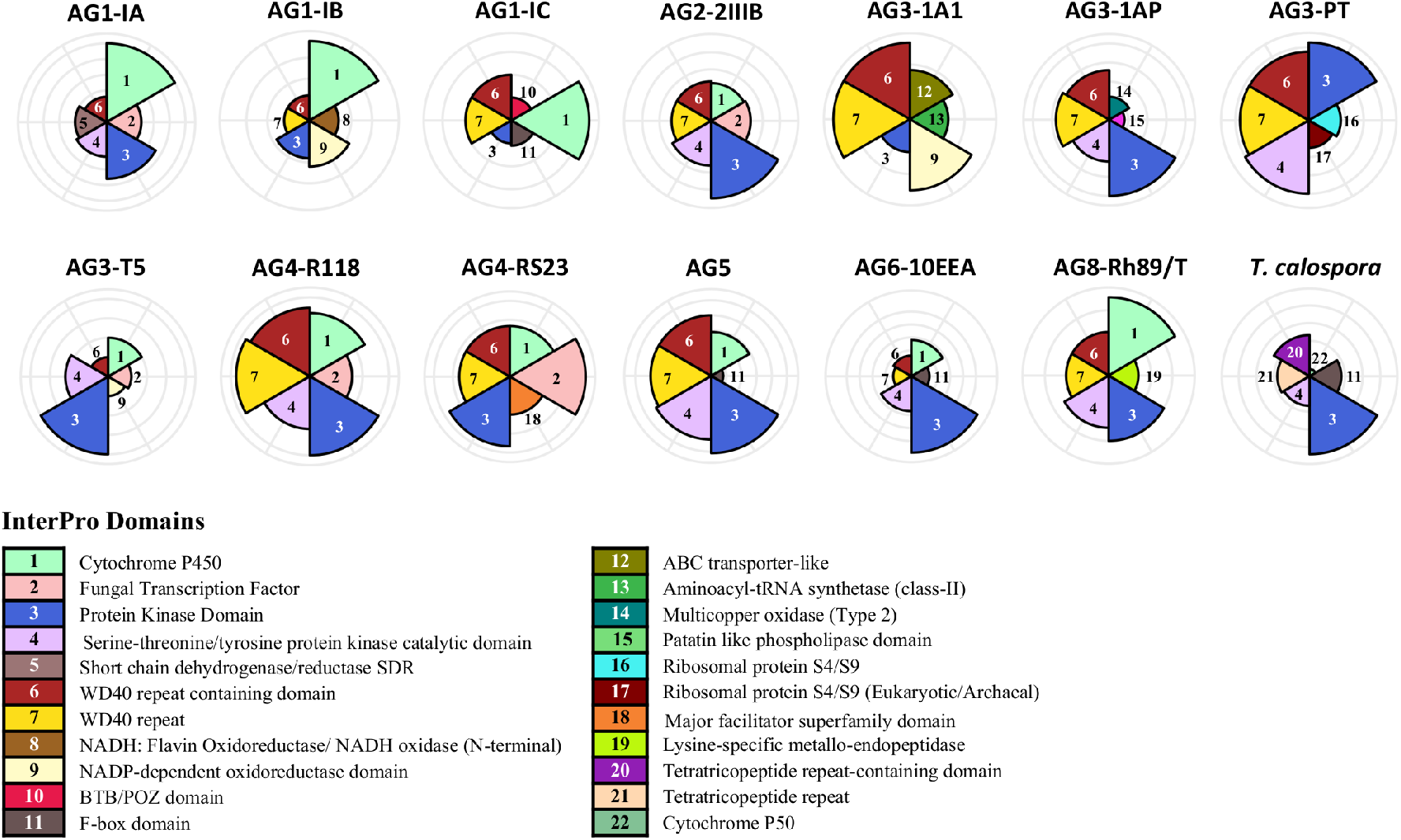
InterPro domain analysis of the unique proteome. In the unique proteome of each fungal isolate, InterPro protein domain families were predicted using InterProScan (Version 5.45-80.0). Only the top 5 most enriched protein families are shown. The number marks the corresponding annotation of InterPro family domain in the circular bar plot.

### The predicted secretome and effector proteins

To facilitate host colonization, plant pathogens secrete proteins to host compartments that modulate morphological changes in the host system and establish fungal infection (68–70). Therefore, we identified the comprehensive set of secreted proteins from all *R. solani* and *T. calospora* genomes. Figure 4A shows the number of secreted proteins identified in each of the given genomes, wherein AG1-IC, AG3-1A1, AG6-10EEA and AG2-2IIIB contains a large number of proteins in the predicted secretome (Supplementary file, sheet 1-2). However, AG1,1C, AG2-2IIIB and AG8 contains a comparatively larger number of isolate-specific secreted proteins (i.e., secreted proteins in the unique proteome), while AG3-1AP, AG3-1A1 and AG3-PT contains comparatively lower number of secreted proteins. Interestingly, InterPro domain analysis of the secreted proteins suggests that the most enriched protein domain in the predicted secretome is “cellulose binding domain – fungal” (Figure 4B) which is essential for the fungal patho-system for the degradation of cellulose and xylans (71). In addition, the secretomes are also enriched with proteins containing “Glycoside Hydrolase Family 61”, “Pectate Lyase” and “multi-copper oxidase family” domains. Most of these protein components include enzymes essential for degradation of the plant host cell wall and breaking down the first line of host defense. We observed that certain families of protein domains are found to be enriched within a few AGs only. For instance, “aspartic peptidase family A1” domain containing proteins, involved in diverse fungal metabolic processes, are mainly enriched in AG2-2IIIB isolate, similarly “lysine-specific metallo-endopeptidase” are enriched in AG3-1AP, AG5 and AG8. The AG4-R118 secretome is significantly enriched with proteins belonging to “Glycoside Hydrolase Family 28” and “Peptidase S8 propeptide-proteinase inhibitor I9” domains, whereas AG4-RS23 secretome is composed of “NodB homology” and “alpha/beta hydrolase fold-1” domains. Taken together, the analysis indicates that each of the given AG secretome is significantly enriched with a unique set of protein families that possibly allows the fungal patho-system to perform a variety of biological functions in different host systems and patho-systems.

**Figure 4.**
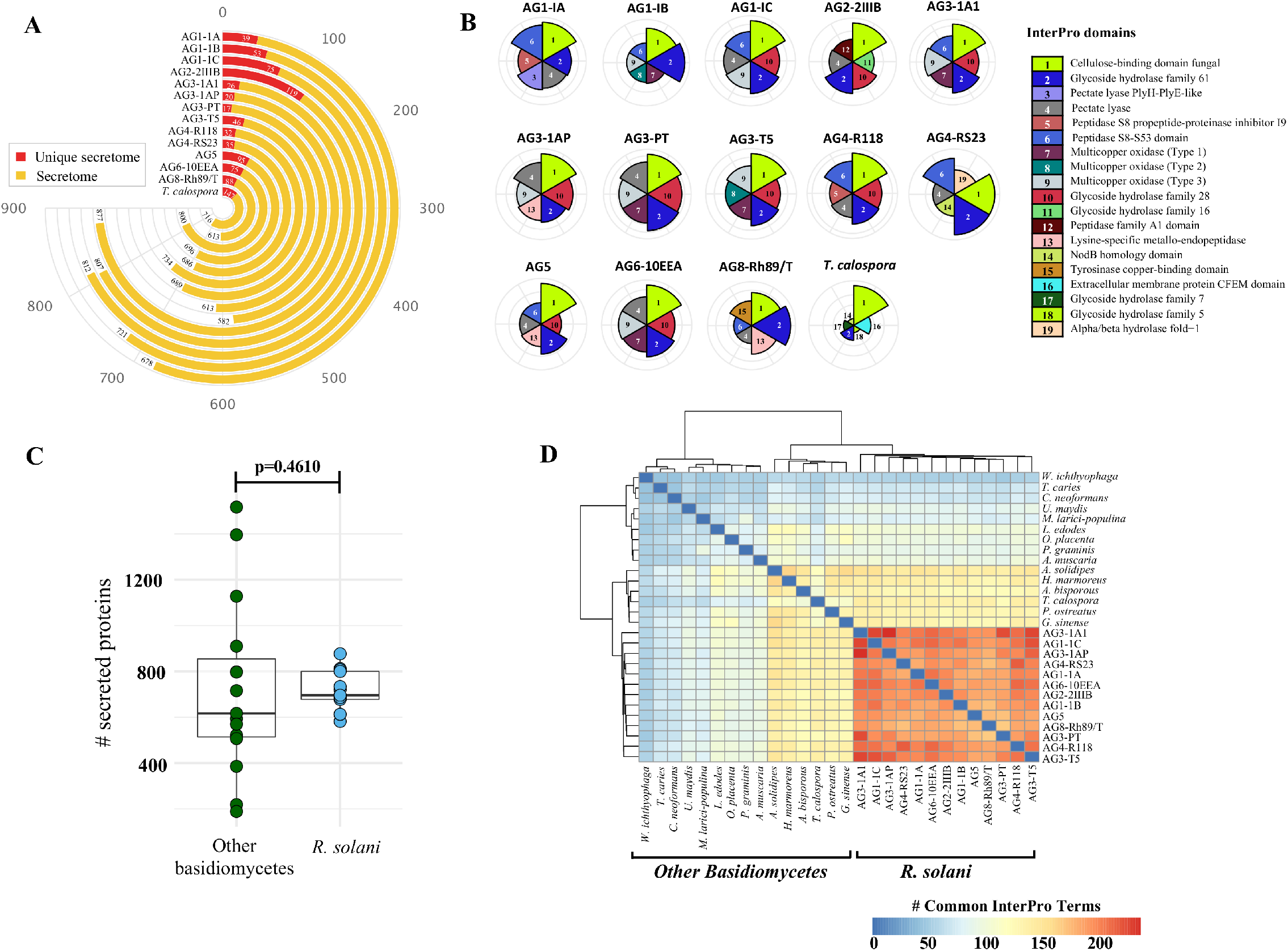
The Secreted Proteins. A. Number of predicted proteins in the secretome of each fungal isolate (highlighted in yellow). The secreted proteins predicted in the unique proteome of each isolate is highlighted in red. B. Comparative analysis of top six highly enriched InterPro domains in the secretome.

Next, to identify the unique and conserved attributes associated with *R solani*, we performed a comparative analysis of the secretome with 14 other fungi (excluding *T. calospora),* which represented the major taxonomic, pathogenic, ecological, and commercially important (edible fungi) groups within the Division Basidiomycota. (Table S4). We hypothesized that small set of functionally important proteins, e.g., secreted proteins, in *R. solani* may have the unique attributes not observed within the other basidiomycetes. Therefore, we predicted the secretome and analyzed the InterPro domains in the secreted proteins of 14 different basidiomycetes and compared with the secretome of *R. solani* AGs. We observed that the number of secreted proteins predicted in *R solani* AGs are not significantly different to the number of secreted proteins in other Basidiomycetes (p=0.0629; Figure 4C). However, the InterPro domains enriched in the secretome of *R. solani* AGs and other basidiomycetes are found to be significantly different. We observed that only a limited number of InterPro terms are shared between *R. solani* AGs and other basidiomycetes, and *R. solani* AGs are functionally closer to each other than other basidiomycetes (Figure 4D), which suggests that *R. solani* secretome have a unique domain profile, which are primarily different from other Basidiomycetes. Overall, we found 565 InterPro terms in the secretome of *R. solani,* whereas in other basidiomycetes (including *T. calospora),* secretomes are enriched with 620 terms in which 283 InterPro terms are common across both the group of species. We observed 282 InterPro terms (50%) uniquely associated with *R. solani,* not observed in the secretome of other basidiomycetes, whereas 337 InterPro terms are only observed in the secretome of other basidiomycetes. The analysis of *R. solani* specific 282 InterPro terms includes several protein domains belonging to diverse functional significance, e.g., “Aspartic peptidase A1 family”, “Cysteine rich secretory protein related” and “Polyscaccaride lyase 8” domains. Among the domains commonly enriched across both *R. solani* isolates and other basidiomycetes, we calculated the fold change of difference of domain occurrence in their secretome and enumerated the proteins domains with significant differences across *R. solani* and other basidiomycetes (Supplementary file, sheet 1-2). Our analysis suggests, high differences in domains frequency wherein, protein with domains like “Pectate lyase”, “Serine amino-peptidase” and “Lysine-specific metallo-endopeptidase” are significantly enriched in *R. solani* secretome. Similarly, proteins with “Hydrophobin” and “Zinc finger ring-type” domains are majorly enriched in other basidiomycetes. We believe that such large number of unique functional domains in the secreted proteome of *R. solani* may be functionally relevant that allows these fungi to survive in diverse array of conditions, and thus should further be investigated experimentally for understanding their role in survival.

Although these plant pathogenic fungi secrete a large number of proteins, only a small proportion of these proteins have been implicated to be effectively associated with fungal-plant interactions, i.e. effector proteins (68–70). Effector proteins can strongly inhibit the activity of host cellular proteases and allow pathogenic fungi to evade host defense mechanisms. Fungal effector proteins are not known for having a conserved family of domains, these proteins typically are of small length (300-400 amino acids) and higher cysteine content (55, 69, 72). Our analysis reveals 75-134 effector proteins predicted in *R. solani* genomes, whereas *T. calospora* contains 136 effector proteins (Figure 5A; supplementary file S1-S7).

**Figure 5.**
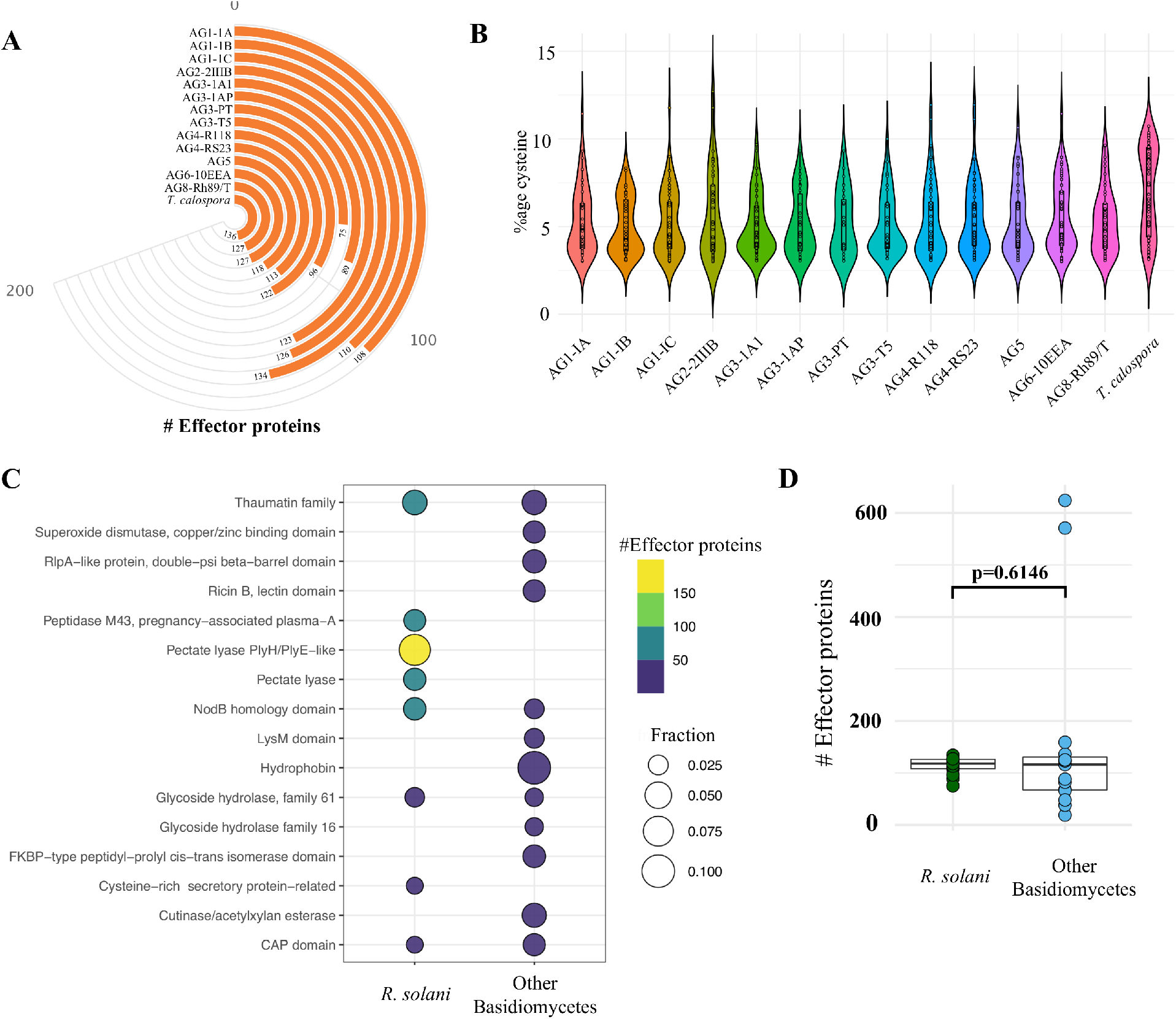
Effector Proteins. A. The number of cysteine rich effector proteins predicted in the predicted secretome of each fungal isolate. B. The proportion of Cysteine observed across all the effectors predicted in each isolate. C. Topmost Enriched InterPro domains in Effector proteins of Rhizoctonia species (not *T. calospora*) and other basidiomycetes (including *T. calospora*). D. The comparative analysis of the distribution of number of effector proteins predicted in *R. solani* AGs as compared to other Basidiomycetes. The p-value is computed using unpaired Wilcoxon-rank sum test.

Isolates from AG1-IC contains the highest number of effector proteins (*n*=134), whereas isolate from AG3-PT contains a small number of effectors (*n*=75). Nevertheless, all the isolates are composed of approximately 100 effector proteins which contain a similar proportion of cysteine residues in the predicted effector proteins (Figure 5B). Next, we investigated the topmost enriched domains among all *R. solani* effector proteins in which “Pectate lyase” is found to be the most enriched effector protein, followed by “thaumatin family” of domain containing proteins (Figure 5C).

In comparison, the analysis of effector proteins in other basidiomycetes suggests that all other basidiomycetes are enriched with similar number of effector proteins (p-value = 0.14; Figure 5C-D; supplementary file; sheet 3-4). Wherein, the effector proteins in *R solani* AGs includes the proteins belonging to 237 InterPro terms, whereas the effector proteins of other basidiomycetes (including *T. calospora)* include proteins enriched with 119 terms. We found 173 terms (72%) are uniquely associated with *R solani* AGs, in which most abundant terms includes IPR001283 (Cystine rich secretory protein related). These unique effectors may play the deciding roles on host recognition and in virulence of necrotrophic *Rhizoctonia* pathogens (73, 74). Moreover, we also observed 55 InterPro terms not observed with *R solani* effector proteins, including Zinc Finger and LysM domain. We also found 64 InterPro terms commonly enriched by both the groups of effector proteins, in which “Pectate lyase” and “Glycoside hydrolase family 28” are mainly associated with *R. solani* AG subgroups effector proteins, whereas “Hydrophobin” is mainly associated with other basidiomycetes. The complete list of secretome, effector proteins, the InterPro domains and associated information are available in supplementary file.

### Carbohydrate-active enzymes

CAZymes are essential for degradation of host plant cells and fungal colonization in the host, and are, thus important for fungal bioactivity (75, 76). Using CAZy (Carbohydrate Active Enzyme database) (77), which contains the classified information of enzymes involved in complex carbohydrate metabolism, we annotated and compared the distribution of CAZymes in all *R. solani* isolates. Overall, *R. solani* isolates are composed of 383-595 high confidence CAZymes, with AG3-1A1 having the largest number of CAZymes (Figure 6A). These predicted CAZymes in *R solani* AGs are mainly distributed across 177 CAZyme families that can be broadly classified into six major classes of enzymes, i.e., Glycoside Hydrolase (GH), Polysaccharide Lyase (PL), Carbohydrate Esterase (CE), Carbohydrate-binding modules (CBM) and redox enzymes with Auxiliary Activities (AA). Our analysis reveals that GH forms the major class of CAZymes in all fungal species, including *T. calospora* (Figure S6 and S7), which hydrolyzes the glycosidic bonds between carbohydrate and non-carbohydrate moieties or between two or more carbohydrate moieties (78). Whereas CBM forms the least abundant class of enzymes enriched in the proteomes of the given isolates. Despite the differences, we observed similar distribution of enzyme count in each class of CAZyme across all the given isolates.

**Figure 6.**
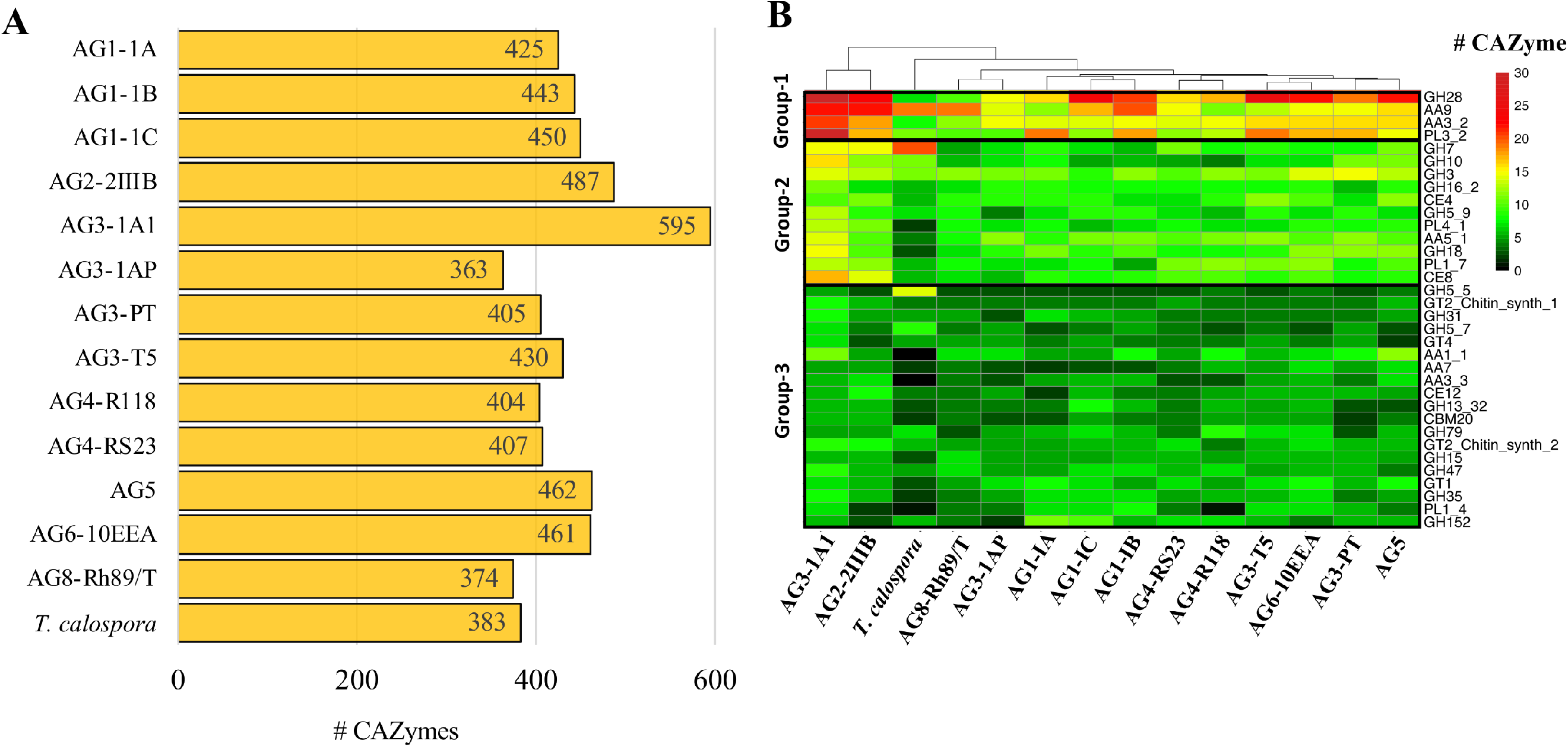
CAZymes. A. The number of carbohydrate metabolizing enzymes (CAZymes) predicted in the proteome of each fungal isolate. B. Heatmap showing the CAZyme conservation across all the *R. solani* AGs and *T. calospora.* Each row represents one CAZy family of proteins, and color is proportional to the number of protein members shared within a given family from the given species (black: no member protein present; red: large number of protein members present). The hierarchical clustering (hclust; method: complete) enumerates the similarities between different fungal isolates based on proteins shared by them across all CAZy families. For simplicity only the CAZyme families enriched in more than 50 enzymes across all proteomes are shown.

We found that among the predicted 177 families, only 34 families are abundant (with total enzyme count > 50 proteins; Figure 6B) across all the given isolates, i.e., *Rhizoctonia* species and *T. calospora.* These 36 families have a distinct abundance profile in each AG, for instance, protein from GH7 family is highly abundant in *T. calospora* as compared to the *R solani* isolates. Similarly, proteins belonging to PL1_4 are not observed in AG4-R118 and *T. calospora.* We have divided these 34 families into three different groups, with respect to their abundance profile in *R. solani* isolates. The Group-1 contains CAZymes belonging to GH28, AA9, PL3_2 and AA3_2 families and form the highly abundant families (total enzyme count >200 proteins) of enzymes in *R Solani* AGs. Similarly, Group-2 contains 11 CAZyme families with enzymes moderately abundant in *R Solani* AGs. Whereas Group-3 contains 19 families with sparsely abundant CAZymes. We observed that in all the three clusters, AG3-1A1 contains the highest number of CAZymes for most the 34 families, and significantly enriched with all the members of Group-1 families. In fact, the clustering analysis highlights the similar profiles of AG3-1A1 and AG2-2IIIB, mainly due to similar distribution of proteins belonging to GH28, AA9, AA3_2 and GH7. In Group-1, although GH28 containing enzymes are abundant in most of the *R. solani* isolates, AG8 contains limited number of enzymes belonging to this family. Similarly, AA9 and PL3_2 families of enzymes are abundant only in 50% of the isolates, and thus may be relevant for a unique set of functions associated with the respective isolates. In Group-2, however, we observed similar distribution of abundance profile across all the isolates, except *T. calospora,* which indicates their probable role in *R. solani* specific function. For examples, CAZymes belonging to AA5_1, GH18 and PL4_1 are enriched in most of the *R. solani* isolates, but not in *T. calospora.* The conserved distribution of CAZymes families in the diverse proteomes of different *R. solani* isolates signifies their essential role in fungal activity. On the other hand, Group-3 CAZymes provide unique and distinct profile to each AG with a limited number of families showing similar abundance profile. Wherein, *T. calospora* is found to be distinctly abundant in CAZymes belonging to GH5_5, not observed with *R. solani* isolates. These results strongly suggested that *R. solani* isolates share a large proportion of carbohydrate degrading enzymes, in which an isolate-specific CAZyme profile can also be observed (mainly from Group-3). To confirm, if the abundance profile is strictly associated with *R. solani* isolates, we performed the comparative analysis with abundance profile of 14 other basidiomycetes. The analysis clearly reveals the distinct CAZymes profile than other Basidiomycetes, in which *R. solani* isolates can be phylogenetically grouped into a different cluster (Figure S8). The analysis highlights the families that uniquely abundant in *R. solani* isolates than other basidiomycetes, e.g., GH28, PL3_2, AA5_1, CE4, GH10, GH62, PL4_1, CE8, PL1_7, PL1_4 and AA7, and as expected, most of these families belong to Group-1 and Group-2 of the previous analysis. Among these families, we observed that PL3_2, GH62 and CE8 families of proteins are distinctly expressed in *R. solani* isolates. In addition, AG3-1A1 is exceptionally abundant in AA9 and GH28, not observed with any other basidiomycetes under investigation. In contrast, AA3_2 (Group-1) is abundant in most of the basidiomycetes, including *R. solani.* In summary, we have shown that members of CAZymes families belonging to Group-1 and Group-2 are abundant in *R. solani* isolates and may also provide them a unique attribute (or functions) not observed with the other basidiomycetes.

### RsolaniDB: a *Rhizoctonia solani* pangenome database and its applications

RDB is a large-scale, integrative repository for hosting the *R. solani* pangenome project with emphasis on supporting data mining and analysis, wherein the genomes and their components can be accessed under three different categories, viz. genomic, ortholog and functional assignment.

#### Genomes

The genomic content includes draft genome sequences of *R. solani* isolates in FASTA format along with the gene level annotation in GFF3 format. The annotation includes prediction of gene boundaries with introns and exons, as well as their locations on contigs or scaffolds. It also includes the predicted transcribed cDNA sequences and translated protein sequences. This information is vital for those users looking for reference genomes and their annotated components for mapping RNAseq reads. The draft genomes and their annotation can also be downloaded and used for downstream local analysis, e.g., variants calling, SNP, eQTLs analysis and other similar genomic analyses with different bioinformatics methods.

#### Orthologs

Using the orthoMCL clustering on the proteomes of 18 *R. solani* (including previously published genome assemblies), protein sequences were compared and clustered into groups of similar sequences. The sequences not part of any of the clusters, i.e., singletons, and unique to respective isolates were categorized as “unique”. Whereas the rest of the proteome was categorized either into “core” or “auxillary” groups of orthoMCL clusters. RDB allows users to retrieve this information for each protein entry and also allows users to retrieve the protein ID of other members of its ortholog cluster family, if any.

#### Functional assignment

This category includes the predicted InterPro protein domains associated with each of the protein entries. RDB also includes GO information associated with each protein, along with PANTHER pathway terms. This information helps in assigning the functional description for each protein entry in the database.

The database is organized to include one unique RDB ID (or entry) for each gene structure, with all of the above associated information. The RDB ID allows users to search the genomic coordinates (intron/exon boundaries) with IGV visualization, sequences and its functional annotation, for each gene in each *R. solani* isolate. All of this information can be retrieved from the database via the “text-based” or “keywords-based” search in an AG-specific manner or from the entire database. Users can also perform blast searches of their own nucleotide or protein sequences to the entire database or can target a given AG. Moreover, users can retrieve the set of sequences in FASTA format, for a given list of RDB IDs. One of the important and unique features of RsolaniDB tools allows users to perform functional or gene-set enrichment analysis of given RDB IDs, e.g., Gene Ontology or pathway analysis. This feature is especially useful for analyzing differentially expressed genes after RNAseq data analysis, as it provides the statistical significance (as *p*-values) of different GO/pathway terms enriched in a given set of differentially expressed genes. As far as we know, this feature is unique to RDB with respect to any other existing *Rhizoctonia* resources. However, it requires the user to use reference genome sequences and the annotation file from RDB database for subjecting into RNAseq data analysis pipeline. As an additional resource, RDB also incorporated previously published (16, 18, 20, 79–81) genome and transcriptome level information in a single platform with an RDB ID format. The database is publicly available to the scientific community, accessible at http://rsolanidb.kaust.edu.sa/RhDB/index.html.

### Discussion

*Rhizoctonia solani* is considered as one of the most destructive and a diverse group of soil-borne plant pathogens causing various diseases on a wide range of economically important crops. It is classified into 13 AGs with distinctive pathogenic host range and responsiveness to disease control measures. For example, AG1, AG2-2IIIB, and AG4 cause diseases mostly on cool-season turfgrasses, whereas AG2-2LP, causing large patch disease, is predominantly seen on warm season turfgrasses (82, 83). Isolates from different AGs also vary in sensitivity to fungicides and no single fungicide is effective against all AGs (84). For example, AG5 isolates are moderately sensitive to pencycuron, while other AGs are highly sensitive to this fungicide (85). Our ability to control this pathogen is hampered by a lack of accurate molecular identification of AGs and its subgroups, and poor understanding of the genetic variation among them. This genetic variation results in differing sensitivity to control measures, as well as the pathogenic and ecological diversity in the population structures of the *R. solani* complex. One of the primary reasons for this limited understanding of the *R. solani* complex is the lack of genetic studies representative of its heterozygous and diverse AGs and sub-groups (13). Until now, draft genome assemblies belonging to only four of the 13 AGs had been reported; viz. AG1-IA (16), AG1-IB (17), AG2-2IIIB (13), AG3-Rhs1AP (18), AG3-PT isolate Ben-3 (19) and AG8 (20). Here we expanded the scope of genetic analysis of the *R. solani* complex by performing comprehensive genome sequencing, assembly, annotation and comparative analysis of 12 *R. solani* isolates. This enabled us to perform pangenome analysis of *R. solani* to 7 AGs (AG1, AG2, AG3, AG4, AG5, AG6, AG8), selected additional sub-groups (AG1-IC, AG3-TB), and a hypovirulent isolate (AG3-1A1). Although heterokarotic and diploid nature of *Rhizoctonia* species are expected to cause the genome assembly challenges (13), in our analysis we observed of a large number of inter-groups syntenic regions and ITS2-based similarities which highlights the high similarities among the given 13 *R. solani* isolates (including AG3-PT). The recognition of conserved ITS2 sequences along with large syntenic regions despite the physiological and taxonomic differences in the given isolates suggests the essentially conserved regions and high quality of the draft genome sequences generated in this study.

Subsequently, to deduce the similarities as well as unique features in the given set of predicted proteomes, we performed a series of comparative analyses that indicated the expected heterogeneity among *R. solani* subgroups with the orchid mycorrhizal fungus *T. calospora* as an outlier. For example, both AG5 and AG2-2IIIB included a large set of unique proteomes as well as secretomes, enriched with InterPro families of proteins that are abundant in these two AGs. Additionally, the proteome of *R. solani* isolates are uniquely and highly enriched with proteins with “pectate lyase” domains, as compared to the other basidiomycetes. Another finding of potential significance is that the highest number of orthoMCL clusters were shared between AG3-1A1 and AG3-1AP, both isolates belonging to the AG3-PT subgroup. Isolate AG3-1A1 is the sector-derived, hypovirulent isolate of the more virulent isolate, AG3-1AP. Intriguingly, AG3-1A1 has been demonstrated to be a successful biocontrol agent of isolate AG3-1AP in the field (86). Competitive niche exclusion is a demonstrated mechanism for biocontrol where the biocontrol agent has a significant overlap in resource utilization with the pathogen and outcompetes the pathogen for these necessary resources (87). A high degree of overlap in gene function is consistent with the mechanism of biocontrol of AG3-1AP in the field by AG3-1A1 through competitive niche exclusion.

The sector-derived, hypovirulent isolate AG3-1A1 however differed from the progenitor isolate AG3-1AP, as well as the other *R. solani* isolates analyzed, in AROM sequences. AROM sequences are known for their conserved profile across fungal species and encode the penta-functional AROM polypeptide that catalyzes five consecutive enzymatic reactions in the prechorismate steps of the shikimate pathway; leading to biosynthesis of the aromatic amino acids tryptophan, tyrosine, and phenylalanine(65). The isolate AG3-1A1 contained two complete penta-functional AROM protein sequences while other isolates contained only one complete sequence or partial AROM sequences. Sectoring as a means of phenotypic plasticity in fungi may take place by genetic mutations, rearrangement of heterokaryotic nuclei, conversion from heterokaryotic to homokaryotic mycelium, exchange of cytoplasmic factors, etc., resulting in changes in morphology, virulence, mating type, sporulation, and ecological adaptations (88). It is possible that the genetic event that led to duplication of the AROM sequences in AG3-1A1 led to hypovirulence. Phenylacetic acid (PAA) has been demonstrated to be a virulence factor in the progenitor isolate, AG3-1AP, and that downregulation of the shikimate pathway occurs in AG3-1A1; resulting in a reduction in production of PAA by AG3-1A1(89). Moreover, possibilities exist that one of the two *arom* genes in AG3-1A1 remains inactive due to methylation, or that the gene duplication is an attempt to compensate for the suppressed shikimate pathway as documented in *Aspergillus nidulans* (65). However, further investigation is necessary to determine if any of those hypotheses is true.

Secretome analysis also revealed several interesting findings that provided unique characteristics to each *R. solani* isolate, e.g., secretome of AG1-1B and AG3-T5 are uniquely and significantly enriched with three different multi-copper oxidases (type 1/2/3), both of which are known to cause foliar diseases. Nevertheless, despite the differences, most of the secretome have similar composition in their significantly enriched protein domains, which mainly includes “Cellulose-binding domain fungal”, “Glycoside hydrolase family 61” and “Pectate lyase”. However, the composition is significantly different with respect to the other basidiomycetes and large number of reported protein families are uniquely associated with multiple *R. solani* isolates. We observed similar finding for the effector proteins, wherein protein containing “Cysteine rich secretory proteins”, “Pectate lyase” and “Thaumatin” are distinctly abundant in *R. solani* isolates, whereas “Hydrophobin” is only abundant in other basidiomycetes. Similarly, the CAZyme analysis highlighted several unique attributes associated with each *R. solani* species especially AG3-1A1 by possessing the CBM1 family of proteins which are linked with degradation of insoluble polysaccharides (90). It was observed that several families of these CAZymes were not present in *T. calospora* which is a symbiotic mycorrhizal fungus and other basidiomycetes, e.g. GH28, PL3_2, AA5_1 and GH10 (91). Overall, data presented in this study are consistent with the hypothesis that AG and sub-groups of *Rhizoctonia* species are highly heterogeneous, each with unique functional genomic properties, while being conserved in their functional regions with respective other groups. However, the unique secretomes, effector and similarly CAZymes profiles of *R. solani* over other basidiomycetes may reflect the ecological and host adaptation strategies, as well as the necrotrophic lifestyle of the former, and call for future research in respective areas to better understand the biology and pathology of the species.

To further propel research with *R. solani* we present our data as the web-resource RsolaniDB (RDB). This web-resource includes detailed information on each *R. solani* isolate, such as the genome properties, predicted transcript/protein sequences, predicted function, and protein orthologues among other AG sub-groups, along with tools for Gene Ontology (GO) and pathway enrichment analysis, orthologs, sequence analysis and IGV visualization of gene models. Also, by adding the previously published genome assemblies and their features, RsolaniDB stands as the universal platform for accessing *R. solani* resources with single identifier format. Since none of the existing Rhizoctonia specific databases host such a large repertoire of genome assemblies and accessory web-tools for functional enrichment analysis of gene set, e.g., differentially expressed genes, RsolaniDB stands as a valuable resource for formulating new hypotheses and understanding the unique or conserved patho-system of *R. solani* AGs and subgroups. The associated gene-set enrichment analysis tool further sets RsolaniDB apart from the existing fungal databases which does not allow the gene enrichment analysis.

Finally, since, each of the *R. solani* AGs or subgroups is characterized by a unique heterogeneous profile, we strongly believe that the presented genome assemblies, annotation and comparative analysis will facilitate mycologists and plant pathologists generating a greater understanding of its biology and ecology, and in developing as well as improving the existing *R. solani* disease management projects, including drug target discovery and design of future diagnostic tools for rapid discrimination of *R. solani* AGs under indoor and outdoor farming environments.

## Supporting information

Supplementary File - Result Tables

## Data availability

All data is publicly available as the error corrected, processed fastq files at European Nucleotide Archive (ENA) at EMBL-EBI under primary accession ID PRJEB39881 (secondary accession: ERP123449) (92, 93). Genome assemblies and corresponding annotations are available at RsolaniDB database (http://rsolanidb.kaust.edu.sa/RhDB/index.html)

## Acknowledgements

The authors thank the members of the Bioscience Core Laboratory (BCL) in KAUST for producing the raw DNA and RNA sequence datasets and Adnan (Ed) Ismaiel (USDA-ARS, SASL, for DNA extraction and fungal culture maintenance). We also thank Drs. Ian Misner and Nadim Alkharouf (Towson University, Towson, MD.) for helping during the initial setting-up of the project.

## Author contributions

A.K., D.K.L. and A.P. conceived the study, interpreted the results and wrote the manuscript; A.K. performed the bioinformatics analysis and developed the computational pipelines and the database; A.R., S.M. and M.N. conducted the molecular experiments, library preparation and sequencing; D.P.R. collected and stored the materials and edited the manuscript; A.P. and D.K.L. supervised the overall project.

## References

1. Yang,G. and Li,C. (2012) General Description of Rhizoctonia Species Complex INTECH Open Access Publisher.

2. Amaradasa,B.S., Horvath,B.J., Lakshman,D.K. and Warnke,S.E. (2013) DNA fingerprinting and anastomosis grouping reveal similar genetic diversity in rhizoctonia species infecting turfgrasses in the transition zone of USA. Mycologia, 105, 1190–1201.

3. Raaijmakers,J.M., Paulitz,T.C., Steinberg,C., Alabouvette,C. and Moёnne-Loccoz,Y. (2009) The rhizosphere: A playground and battlefield for soilborne pathogens and beneficial microorganisms. Plant Soil, 321, 341–361.

4. Gónzalez,D., Rodriguez-Carres,M., Boekhout,T., Stalpers,J., Kuramae,E.E., Nakatani,A.K., Vilgalys,R. and Cubeta,M.A. (2016) Phylogenetic relationships of Rhizoctonia fungi within the Cantharellales. Fungal Biol., 120, 603–619.

5. Keijer,J., Korsman,M.G., Dullemans,A.M., Houterman,P.M., De Bree,J. and Van Silfhout,C.H. (1997) In vitro analysis of host plant specificity in Rhizoctonia solani. Plant Pathol., 46, 659–669.

6. Foley,R.C., Gleason,C.A., Anderson,J.P., Hamann,T. and Singh,K.B. (2013) Genetic and Genomic Analysis of Rhizoctonia solani Interactions with Arabidopsis; Evidence of Resistance Mediated through NADPH Oxidases. PLoS One, 8, e56814.

7. Gonzalez,D., Carling,D.E., Kuninaga,S., Vilgalys,R. and Cubeta,M.A. (2001) Ribosomal DNA systematics of Ceratobasidium and Thanatephorus with Rhizoctonia anamorphs. Mycologia, 93, 1138–1150.

8. Hane,J.K., Anderson,J.P., Williams,A.H., Sperschneider,J. and Singh,K.B. (2014) Genome sequencing and comparative genomics of the broad host-range pathogen Rhizoctonia solani AG8. PLoS Genet., 10, e1004281.

9. Hossain,M.K., Tze,O.S., Nadarajah,K., Jena,K., Bhuiyan,M.A.R. and Ratnam,W. (2014) Identification and validation of sheath blight resistance in rice (Oryza sativa L.) cultivars against Rhizoctonia solani. Can. J. Plant Pathol., 36, 482–490.

10. Copley,T., Bayen,S. and Jabaji,S. (2017) Biochar Amendment Modifies Expression of Soybean and Rhizoctonia solani Genes Leading to Increased Severity of Rhizoctonia Foliar Blight. Front. Plant Sci., 8, 221.

11. Anderson,J.P., Hane,J.K., Stoll,T., Pain,N., Hastie,M.L., Kaur,P., Hoogland,C., Gorman,J.J. and Singh,K.B. (2016) Proteomic analysis of rhizoctonia solani identifies infection-specific, redox associated proteins and insight into adaptation to different plant hosts. Mol. Cell. Proteomics, 15, 1188–1203.

12. Lakshman,D.K., Roberts,D.P., Garrett,W.M., Natarajan,S.S., Darwish,O., Alkharouf,N., Pain,A., Khan,F., Jambhulkar,P.P. and Mitra,A. (2016) Proteomic Investigation of Rhizoctonia solani AG 4 Identifies Secretome and Mycelial Proteins with Roles in Plant Cell Wall Degradation and Virulence. J. Agric. Food Chem., 64, 3101–3110.

13. Wibberg,D., Andersson,L., Tzelepis,G., Rupp,O., Blom,J., Jelonek,L., Pühler,A., Fogelqvist,J., Varrelmann,M., Schlüter,A., et al. (2016) Genome analysis of the sugar beet pathogen Rhizoctonia solani AG2-2IIIB revealed high numbers in secreted proteins and cell wall degrading enzymes. BMC Genomics, 17, 245.

14. Zhang,J., Chen,L., Fu,C., Wang,L., Liu,H., Cheng,Y., Li,S., Deng,Q., Wang,S., Zhu,J., et al. (2017) Comparative transcriptome analyses of gene expression changes triggered by Rhizoctonia solani AG1 IA infection in resistant and susceptible rice varieties. Front. Plant Sci., 8, 1422.

15. Shu,C., Zhao,M., Anderson,J.P., Garg,G., Singh,K.B., Zheng,W., Wang,C., Yang,M. and Zhou,E. (2019) Transcriptome analysis reveals molecular mechanisms of sclerotial development in the rice sheath blight pathogen Rhizoctonia solani AG1-IA. Funct. Integr. Genomics, 19, 743–758.

16. Nadarajah,K., Razali,N.M., Cheah,B.H., Sahruna,N.S., Ismail,I., Tathode,M. and Bankar,K. (2017) Draft genome sequence of Rhizoctonia solani anastomosis group 1 subgroup 1A strain 1802/KB isolated from rice. Genome Announc., 5.

17. Wibberg,D., Rupp,O., Blom,J., Jelonek,L., Kröber,M., Verwaaijen,B., Goesmann,A., Albaum,S., Grosch,R., Pühler,A., et al. (2015) Development of a Rhizoctonia solani AG1-IB Specific Gene Model Enables Comparative Genome Analyses between Phytopathogenic R. solani AG1-IA, AG1-IB, AG3 and AG8 Isolates. PLoS One, 10.

18. Cubeta,M.A., Thomas,E., Dean,R.A., Jabaji,S., Neate,S.M., Tavantzis,S., Toda,T., Vilgalys,R., Bharathan,N., Fedorova-Abrams,N., et al. (2014) Draft Genome Sequence of the Plant-Pathogenic Soil Fungus Rhizoctonia solani Anastomosis Group 3 Strain Rhs1AP. Genome Announc., 2.

19. Wibberg,D., Genzel,F., Verwaaijen,B., Blom,J., Rupp,O., Goesmann,A., Zrenner,R., Grosch,R., Pühler,A. and Schlüter,A. (2017) Draft genome sequence of the potato pathogen Rhizoctonia solani AG3-PT isolate Ben3. Arch. Microbiol., 199, 1065–1068.

20. Hane,J.K., Anderson,J.P., Williams,A.H., Sperschneider,J. and Singh,K.B. (2014) Genome Sequencing and Comparative Genomics of the Broad Host-Range Pathogen Rhizoctonia solani AG8. PLoS Genet., 10.

21. Bills,G.F., Singleton,L.L., Mihail,J.D. and Rush,C.M. (1993) Methods for Research on Soilborne Phytopathogenic Fungi The American Phytopathological Society,.

22. Carlson,J.E., Tulsieram,L.K., Glaubitz,J.C., Luk,V.W.K., Kauffeldt,C. and Rutledge,R. (1991) Segregation of random amplified DNA markers in F1 progeny of conifers. Theor. Appl. Genet., 10.1007/BF00226251.

23. Sayers,E.W., Beck,J., Brister,J.R., Bolton,E.E., Canese,K., Comeau,D.C., Funk,K., Ketter,A., Kim,S., Kimchi,A., et al. (2020) Database resources of the National Center for Biotechnology Information. Nucleic Acids Res., 10.1093/nar/gkz899.

24. Bolger,A.M., Lohse,M. and Usadel,B. (2014) Trimmomatic: A flexible trimmer for Illumina sequence data. Bioinformatics, 30, 2114–2120.

25. Simon Andrews (2020) Babraham Bioinformatics - FastQC A Quality Control tool for High Throughput Sequence Data. Soil, 5, 47–81.

26. Marçais,G. and Kingsford,C. (2011) A fast, lock-free approach for efficient parallel counting of occurrences of k-mers. Bioinformatics, 27, 764–770.

27. Grabherr,M.G., Haas,B.J., Yassour,M., Levin,J.Z., Thompson,D.A., Amit,I., Adiconis,X., Fan,L., Raychowdhury,R., Zeng,Q., et al. (2011) Full-length transcriptome assembly from RNA-Seq data without a reference genome. Nat. Biotechnol., 29, 644–652.

28. Bankevich,A., Nurk,S., Antipov,D., Gurevich,A.A., Dvorkin,M., Kulikov,A.S., Lesin,V.M., Nikolenko,S.I., Pham,S., Prjibelski,A.D., et al. (2012) SPAdes: A new genome assembly algorithm and its applications to single-cell sequencing. J. Comput. Biol., 19, 455–477.

29. Gurevich,A., Saveliev,V., Vyahhi,N. and Tesler,G. (2013) QUAST: Quality assessment tool for genome assemblies. Bioinformatics, 29, 1072–1075.

30. Boetzer,M., Henkel,C. V., Jansen,H.J., Butler,D. and Pirovano,W. (2011) Scaffolding pre-assembled contigs using SSPACE. Bioinformatics, 10.1093/bioinformatics/btq683.

31. Luo,R., Liu,B., Xie,Y., Li,Z., Huang,W., Yuan,J., He,G., Chen,Y., Pan,Q., Liu,Y., et al. (2012) SOAPdenovo2: An empirically improved memory-efficient short-read de novo assembler. Gigascience, 1, 18.

32. Song,L., Shankar,D.S. and Florea,L. (2016) Rascaf: Improving Genome Assembly with RNA Sequencing Data. Plant Genome, 9, plantgenome2016.03.0027.

33. Seppey,M., Manni,M. and Zdobnov,E.M. (2019) BUSCO: Assessing genome assembly and annotation completeness. In Methods in Molecular Biology. Humana Press Inc., Vol. 1962, pp. 227–245.

34. Bengtsson-Palme,J., Ryberg,M., Hartmann,M., Branco,S., Wang,Z., Godhe,A., De Wit,P., Sánchez-García,M., Ebersberger,I., de Sousa,F., et al. (2013) Improved software detection and extraction of ITS1 and ITS2 from ribosomal ITS sequences of fungi and other eukaryotes for analysis of environmental sequencing data. Methods Ecol. Evol., 4, 914–919.

35. Kumar,S., Stecher,G., Li,M., Knyaz,C. and Tamura,K. (2018) MEGA X: Molecular evolutionary genetics analysis across computing platforms. Mol. Biol. Evol., 10.1093/molbev/msy096.

36. Rédei,G.P. (2008) CLUSTAL W (improving the sensitivity of progressive multiple sequence alignment through sequence weighting, position-specific gap penalties and weight matrix choice). In Encyclopedia of Genetics, Genomics, Proteomics and Informatics.

37. Jovanovic,N. and Mikheyev,A.S. (2019) Interactive web-based visualization and sharing of phylogenetic trees using phylogeny.IO. Nucleic Acids Res., 47, W266–W269.

38. Huerta-Cepas,J., Serra,F. and Bork,P. (2016) ETE 3: Reconstruction, Analysis, and Visualization of Phylogenomic Data. Mol. Biol. Evol., 33, 1635–1638.

39. Pryszcz,L.P. and Gabaldón,T. (2016) Redundans: An assembly pipeline for highly heterozygous genomes. Nucleic Acids Res., 44, e113.

40. Camacho,C., Coulouris,G., Avagyan,V., Ma,N., Papadopoulos,J., Bealer,K. and Madden,T.L. (2009) BLAST+: Architecture and applications. BMC Bioinformatics, 10, 421.

41. Cantarel,B.L., Korf,I., Robb,S.M.C., Parra,G., Ross,E., Moore,B., Holt,C., Alvarado,A.S. and Yandell,M. (2008) MAKER: An easy-to-use annotation pipeline designed for emerging model organism genomes. Genome Res., 18, 188–196.

42. Tarailo-Graovac,M. and Chen,N. (2009) Using RepeatMasker to identify repetitive elements in genomic sequences. Curr. Protoc. Bioinforma., 10.1002/0471250953.bi0410s25.

43. Bateman,A., Martin,M.J., O’Donovan,C., Magrane,M., Alpi,E., Antunes,R., Bely,B., Bingley,M., Bonilla,C., Britto,R., et al. (2017) UniProt: The universal protein knowledgebase. Nucleic Acids Res., 10.1093/nar/gkw1099.

44. Mi,H. and Thomas,P. (2009) PANTHER pathway: an ontology-based pathway database coupled with data analysis tools. Methods Mol. Biol., 563, 123–140.

45. Ashburner,M., Ball,C.A., Blake,J.A., Botstein,D., Butler,H., Cherry,J.M., Davis,A.P., Dolinski,K., Dwight,S.S., Eppig,J.T., et al. (2000) Gene ontology: Tool for the unification of biology. Nat. Genet., 25, 25–29.

46. Quevillon,E., Silventoinen,V., Pillai,S., Harte,N., Mulder,N., Apweiler,R. and Lopez,R. (2005) InterProScan: Protein domains identifier. Nucleic Acids Res., 33.

47. Hulo,N. (2004) Recent improvements to the PROSITE database. Nucleic Acids Res., 10.1093/nar/gkh044.

48. Marchler-Bauer,A., Zheng,C., Chitsaz,F., Derbyshire,M.K., Geer,L.Y., Geer,R.C., Gonzales,N.R., Gwadz,M., Hurwitz,D.I., Lanczycki,C.J., et al. (2013) CDD: Conserved domains and protein three-dimensional structure. Nucleic Acids Res., 10.1093/nar/gks1243.

49. Finn,R.D., Bateman,A., Clements,J., Coggill,P., Eberhardt,R.Y., Eddy,S.R., Heger,A., Hetherington,K., Holm,L., Mistry,J., et al. (2014) Pfam: The protein families database. Nucleic Acids Res., 10.1093/nar/gkt1223.

50. Haft,D.H., Selengut,J.D. and White,O. (2003) The TIGRFAMs database of protein families. Nucleic Acids Res., 10.1093/nar/gkg128.

51. Potter,S.C., Luciani,A., Eddy,S.R., Park,Y., Lopez,R. and Finn,R.D. (2018) HMMER web server: 2018 update. Nucleic Acids Res., 46, W200–W204.

52. Almagro Armenteros,J.J., Tsirigos,K.D., Sønderby,C.K., Petersen,T.N., Winther,O., Brunak,S., von Heijne,G. and Nielsen,H. (2019) SignalP 5.0 improves signal peptide predictions using deep neural networks. Nat. Biotechnol., 37, 420–423.

53. Käll,L., Krogh,A. and Sonnhammer,E.L.L. (2007) Advantages of combined transmembrane topology and signal peptide prediction-the Phobius web server. Nucleic Acids Res., 35.

54. Emanuelsson,O., Brunak,S., von Heijne,G. and Nielsen,H. (2007) Locating proteins in the cell using TargetP, SignalP and related tools. Nat. Protoc., 2, 953–971.

55. Sperschneider,J., Gardiner,D.M., Dodds,P.N., Tini,F., Covarelli,L., Singh,K.B., Manners,J.M. and Taylor,J.M. (2016) EffectorP: Predicting fungal effector proteins from secretomes using machine learning. New Phytol., 210, 743–761.

56. Zhang,H., Yohe,T., Huang,L., Entwistle,S., Wu,P., Yang,Z., Busk,P.K., Xu,Y. and Yin,Y. (2018) DbCAN2: A meta server for automated carbohydrate-active enzyme annotation. Nucleic Acids Res., 46, W95–W101.

57. Farrer,R.A. (2017) Synima: A Synteny imaging tool for annotated genome assemblies. BMC Bioinformatics, 18, 507.

58. Li,L., Stoeckert,C.J. and Roos,D.S. (2003) OrthoMCL: Identification of ortholog groups for eukaryotic genomes. Genome Res., 13, 2178–2189.

59. Yu,Y., Ouyang,Y. and Yao,W. (2018) ShinyCircos: An R/Shiny application for interactive creation of Circos plot. Bioinformatics, 10.1093/bioinformatics/btx763.

60. Krzywinski,M., Schein,J., Birol,I., Connors,J., Gascoyne,R., Horsman,D., Jones,S.J. and Marra,M.A. (2009) Circos: An information aesthetic for comparative genomics. Genome Res., 10.1101/gr.092759.109.

61. Alexa,A. and Rahnenführer,J. (2009) Gene set enrichment analysis with topGO. Bioconductor Improv, 27.

62. Marçais,G., Delcher,A.L., Phillippy,A.M., Coston,R., Salzberg,S.L. and Zimin,A. (2018) MUMmer4: A fast and versatile genome alignment system. PLoS Comput. Biol., 14.

63. Cubeta,M.A., Thomas,E., Dean,R.A., Jabaji,S., Neate,S.M., Tavantzis,S., Toda,T., Vilgalys,R., Bharathan,N., Fedorova-Abrams,N., et al. (2014) Draft genome sequence of the plant-pathogenic soil fungus Rhizoctonia solani anastomosis group 3 strain Rhs1AP. Genome Announc., 10.1128/genomeA.01072-14.

64. Lakshman,D.K., Liu,C., Mishra,P.K. and Tavantzis,S. (2006) Characterization of the arom gene in Rhizoctonia solani, and transcription patterns under stable and induced hypovirulence conditions. Curr. Genet., 10.1007/s00294-005-0005-6.

65. Lamb,H.K., Van Den Hombergh,J.P.T.W., Newton,G.H., Moore,J.D., Roberts,C.F. and Hawkins,A.R. (1992) Differential flux through the quinate and shikimate pathways: Implications for the channelling hypothesis. Biochem. J., 10.1042/bj2840181.

66. Lakshman,D.K., Jian,J. and Tavantzis,S.M. (1998) A double-stranded RNA element from a hypovirulent strain of Rhizoctonia solani occurs in DNA form and is genetically related to the pentafunctional AROM protein of the shikimate pathway. Proc. Natl. Acad. Sci. U. S. A., 95, 6425–6429.

67. Črešnar,B. and Petrič,Š. (2011) Cytochrome P450 enzymes in the fungal kingdom. Biochim. Biophys. Acta - Proteins Proteomics, 10.1016/j.bbapap.2010.06.020.

68. Kim,K.T., Jeon,J., Choi,J., Cheong,K., Song,H., Choi,G., Kang,S. and Lee,Y.H. (2016) Kingdom-wide analysis of fungal small secreted proteins (SSPs) reveals their potential role in host association. Front. Plant Sci., 7.

69. McCotter,S.W., Horianopoulos,L.C. and Kronstad,J.W. (2016) Regulation of the fungal secretome. Curr. Genet., 62, 533–545.

70. Li,T., Wu,Y., Wang,Y., Gao,H., Gupta,V.K., Duan,X., Qu,H. and Jiang,Y. (2019) Secretome profiling reveals virulence-associated proteins of Fusarium proliferatum during interaction with banana fruit. Biomolecules, 9.

71. Linder,M., Lindeberg,G., Reinikainen,T., Teeri,T.T. and Pettersson,G. (1995) The difference in affinity between two fungal cellulose-binding domains is dominated by a single amino acid substitution. FEBS Lett., 10.1016/0014-5793(95)00961-8.

72. Stergiopoulos,I. and de Wit,P.J.G.M. (2009) Fungal Effector Proteins. Annu. Rev. Phytopathol., 47, 233–263.

73. Wei,M., Wang,A., Liu,Y., Ma,L., Niu,X. and Zheng,A. (2020) Identification of the Novel Effector RsIA_NP8 in Rhizoctonia solani AG1 IA That Induces Cell Death and Triggers Defense Responses in Non-Host Plants. Front. Microbiol., 10.3389/fmicb.2020.01115.

74. Yamamoto,N., Wang,Y., Lin,R., Liang,Y., Liu,Y., Zhu,J., Wang,L., Wang,S., Liu,H., Deng,Q., et al. (2019) Integrative transcriptome analysis discloses the molecular basis of a heterogeneous fungal phytopathogen complex, Rhizoctonia solani AG-1 subgroups. Sci. Rep., 9.

75. Kameshwar,A.K.S., Ramos,L.P. and Qin,W. (2019) CAZymes-based ranking of fungi (CBRF): an interactive web database for identifying fungi with extrinsic plant biomass degrading abilities. Bioresour. Bioprocess., 10.1186/s40643-019-0286-0.

76. Barrett,K., Jensen,K., Meyer,A.S., Frisvad,J.C. and Lange,L. (2020) Fungal secretome profile categorization of CAZymes by function and family corresponds to fungal phylogeny and taxonomy: Example Aspergillus and Penicillium. Sci. Rep., 10.1038/s41598-020-61907-1.

77. Lombard,V., Golaconda Ramulu,H., Drula,E., Coutinho,P.M. and Henrissat,B. (2014) The carbohydrate-active enzymes database (CAZy) in 2013. Nucleic Acids Res., 10.1093/nar/gkt1178.

78. Henrissat,B. (1991) A classification of glycosyl hydrolases based on amino acid sequence similarities. Biochem. J., 10.1042/bj2800309.

79. Wibberg,D., Genzel,F., Verwaaijen,B., Blom,J., Rupp,O., Goesmann,A., Zrenner,R., Grosch,R., Pühler,A. and Schlüter,A. (2017) Draft genome sequence of the potato pathogen Rhizoctonia solani AG3-PT isolate Ben3. Arch. Microbiol., 199, 1065–1068.

80. Wibberg,D., Andersson,L., Rupp,O., Goesmann,A., Pühler,A., Varrelmann,M., Dixelius,C. and Schlüter,A. (2016) Draft genome sequence of the sugar beet pathogen Rhizoctonia solani AG2-2IIIB strain BBA69670. J. Biotechnol., 10.1016/j.jbiotec.2016.02.001.

81. Wibberg,D., Rupp,O., Jelonek,L., Kröber,M., Verwaaijen,B., Blom,J., Winkler,A., Goesmann,A., Grosch,R., Pühler,A., et al. (2015) Improved genome sequence of the phytopathogenic fungus Rhizoctonia solani AG1-IB 7/3/14 as established by deep mate-pair sequencing on the MiSeq (Illumina) system. J. Biotechnol., 10.1016/j.jbiotec.2015.03.005.

82. Richard W. Smiley, Peter H. Dernoeden, and B.B.C. (2005) Compendium of Turfgrass Diseases, Third Edition.

83. Burpee,L.L. and Martin,S.B. (1996) Biology of Turfgrass Diseases Incited by Rhizoctonia Species. In Rhizoctonia Species: Taxonomy, Molecular Biology, Ecology, Pathology and Disease Control.

84. Amaradasa,B.S., Lakshman,D., Mccall,D.S. and Horvath,B.J. (2014) In Vitro Fungicide Sensitivity of Rhizoctonia and Waitea Isolates Collected from Turfgrasses 1.

85. Campion,C., Chatot,C., Perraton,B. and Andrivon,D. (2003) Anastomosis groups, pathogenicity and sensitivity to fungicides of Rhizoctonia solani isolates collected on potato crops in France. Eur. J. Plant Pathol., 10.1023/B:EJPP.0000003829.83671.8f.

86. Bernard,E., Larkin,R.P., Tavantzis,S., Erich,M.S., Alyokhin,A., Sewell,G., Lannan,A. and Gross,S.D. (2012) Compost, rapeseed rotation, and biocontrol agents significantly impact soil microbial communities in organic and conventional potato production systems. Appl. Soil Ecol., 10.1016/j.apsoil.2011.10.002.

87. Roberts,D.P. and Kobayashi,D.Y. (2011) Impact of Spatial Heterogeneity Within Spermosphere and Rhizosphere Environments on Performance of Bacterial Biological Control Agents. In Bacteria in Agrobiology: Crop Ecosystems.

88. Roper,M., Simonin,A., Hickey,P.C., Leeder,A. and Glass,N.L. (2013) Nuclear dynamics in a fungal chimera. Proc. Natl. Acad. Sci. U. S. A., 10.1073/pnas.1220842110.

89. Liu,C., Lakshman,D.K. and Tavantzis,S.M. (2003) Quinic acid induces hypovirulence and expression of a hypovirulence-associated double-stranded RNA in Rhizoctonia solani. Curr. Genet., 10.1007/s00294-003-0375-6.

90. Van Bueren,A.L., Morland,C., Gilbert,H.J. and Boraston,A.B. (2005) Family 6 carbohydrate binding modules recognize the non-reducing end of β-1,3-linked glucans by presenting a unique ligand binding surface. J. Biol. Chem., 10.1074/jbc.M410113200.

91. Fochi,V., Chitarra,W., Kohler,A., Voyron,S., Singan,V.R., Lindquist,E.A., Barry,K.W., Girlanda,M., Grigoriev,I. V., Martin,F., et al. (2017) Fungal and plant gene expression in the Tulasnella calospora–Serapias vomeracea symbiosis provides clues about nitrogen pathways in orchid mycorrhizas. New Phytol., 10.1111/nph.14279.

92. Harrison,P.W., Alako,B., Amid,C., Cerdeño-Tárraga,A., Cleland,I., Holt,S., Hussein,A., Jayathilaka,S., Kay,S., Keane,T., et al. (2019) The European Nucleotide Archive in 2018. Nucleic Acids Res., 10.1093/nar/gky1078.

93. Leinonen,R., Akhtar,R., Birney,E., Bower,L., Cerdeno-Tárraga,A., Cheng,Y., Cleland,I., Faruque,N., Goodgame,N., Gibson,R., et al. (2011) The European nucleotide archive. Nucleic Acids Res., 10.1093/nar/gkq967.

