## Supplementary material for "Pangenome analysis of the soil-borne fungal phytopathogen *Rhizoctonia solani* and development of a comprehensive web resource: RsolaniDB": Supplemetary Info- Text, Tables and Figures

**Supplementary information**

#### Supplementary tables

**Table S1.** Details of *Rhizoctonia solani* cultures used in the genomic investigation

| Isolate number | AGs | AG subgroups | Isolate name | Plant Host | Reference | Obtained from |
|---|---|---|---|---|---|---|
| 1 | AG1 | AG1-IA | BM1 | Unknown | *in-lab* isolate | BM |
| 2 | | AG1-IB | BM2 | Unknown | (1) | BM |
| 3 | | AG1-IC* | BM3 | Unknown | (1) | BM |
| 4 | AG2 | AG2-2IIIB | Rh-146 | Bentgrass, Georgia, USA | (1) | LB |
| 5 | AG3 | AG3-PT | Rhs1AP or 1AP | Potato, Maine, USA (This isolate was sequenced by Cubeta et al and named as Rhs 1AP) | (2, 3) | |
| 6 | | | Rhs1A1* or 1A1 | Sectored, hypovirulent isolate of Rhs 1AP | (2) | *In lab* isolate |
| 7 | | AG3-TB* | AG3-T5 or T5 | Tobacco, North Carolina, USA | (4) | MC |
| 8 | AG4* | HG-I | R118-11 | A basidiospore isolate of parent isolate R118 isolated from soil, Maryland, USA | (5) | American Type Culture Collection, CN 16118 |
| 9 | | | RS23A or Rs23 | Soil, Maryland, USA | (6, 7) | *In lab* isolate |
| 10 | AG5* | N/D | 63/T | Unknown | (1) | LB |
| 11 | AG6* | N/D | 10EEA | Sweet gum, North Carolina, USA | (1) | MC |
| 12 | AG8 | N/D | Rh89/T | Oat root, Scotland | (1) | LB |

N/D = Sub-groups not identified or does not exist;
BM: Bruce Martin, Clemson University, South Carolina;
LB: Lee Burpee, University of Georgia;
MC: Marc Cubeta, North Carolina State University.
* first genome sequences reported in this study for a marked AG, a sub-group, or a sector-derived hypovirulent isolate.

19

**Table S2**: Summary of genome libraries and sequencing reads.

| AG subgroups | Isolate number | Library Type | Read Count[#] | GC % | Mean read length | RNAseq available |
|---|---|---|---|---|---|---|
| AG1-IA | BM1 | Paired end | ~97M | 45 | 86 | Yes |
| AG1-IB | BM2 | Paired end | ~52M | 45 | 80 | - |
| | | Paired end | ~65M | 46 | 80 | |
| AG1-IC | BM3 | Paired end | ~39M | 46 | 76 | - |
| | | Mate-pair | ~33M | 45 | 76 | |
| AG2-2IIIB | Rh-146 | Paired end | ~157M | 45 | 85 | Yes |
| | | Paired end | ~41M | 46 | 82 | |
| | | Single end | ~41M | 46 | 89 | |
| AG3-1A1 | Rhs1A1 | Paired end | ~105M | 44 | 76 | Yes |
| | | Paired end | ~37M | 47 | 82 | |
| | | Mate-pair | ~21M | 47 | 69 | |
| AG3-1AP | Rhs1AP | Paired end | ~97M | 45 | 85 | - |
| AG3-T5 | * | Paired end | ~103M | 45 | 79 | Yes |
| | | Paired end | ~43M | 46 | 71 | |
| | | Mate-pair | ~26M | 47 | 81 | |
| AG4-R118 | * | Paired end | ~44M | 46 | 84 | Yes |
| | | Paired end | ~37M | 46 | 77 | |
| | | Mate-pair | ~25M | 47 | 77 | |
| AG4-RS23 | * | Paired end | ~130M | 46 | 84 | - |
| AG5 | 63/T | Paired end | ~38M | 47 | 76 | Yes |
| | | Mate-pair | ~27M | 48 | 77 | |
| AG6-10EEA | * | Paired end | ~42M | 47 | 76 | - |
| | | Mate-pair | ~42M | 46 | 75 | |
| AG8 | Rh89/T | Paired end | ~22M | 46 | 80 | - |
| | | Paired end | ~48M | 46 | 71 | |
| | | Paired end | ~37M | 45 | 76 | |
| | | Mate-pair | ~44M | 45 | 76 | |

### in millions; QC, trimmed and error corrected reads

* Not available

| **Table S3: Assembly properties of genomes** reported in this study | | | | | | | | |
|---|---|---|---|---|---|---|---|---|
| | Genome Length | GC (%) | # contigs | N50 | # N's per 100 kbp | Largest contig length | BUSCO | Repetitive genome (%age) |
| AG1-1A | 46,986,790 | 47.47 | 9,145 | 13,594 | 10.92 | 162,406 | 89.30% | 1.47 |
| AG1-1B | 34,314,604 | 48.34 | 1,833 | 69,438 | 3.86 | 607,833 | 95.80% | 0.68 |
| AG1-1C | 33,636,943 | 48.07 | 1,221 | 100,597 | 800.74 | 921,812 | 93.40% | 0.56 |
| AG2-2IIIB | 57,816,428 | 48 | 10,766 | 11,419 | 5.53 | 76,418 | 92.10% | 0.82 |
| AG3-1A1 | 71,708,704 | 49.05 | 7,116 | 29,147 | 579.62 | 485,982 | 96.50% | 0.72 |
| AG3-1AP | 46,445,294 | 48.23 | 10,166 | 8,499 | 7.87 | 213,864 | 90.30% | 0.78 |
| AG3-T5 | 35,199,394 | 48.68 | 678 | 196,133 | 1629.7 | 79,7852 | 95.20% | 0.75 |
| AG4-R118 | 38,232,519 | 48.45 | 1,860 | 52,568 | 597.37 | 638,355 | 91.30% | 0.65 |
| AG4-RS23 | 42,733,718 | 48.22 | 4,504 | 19,406 | 47.48 | 232,254 | 89.60% | 0.88 |
| AG5 | 47,761,425 | 49.01 | 6,674 | 12,763 | 1961.37 | 85,396 | 88.70% | 0.58 |
| AG6-10EEA | 47,563,091 | 49.07 | 4,274 | 25,715 | 1465.94 | 373,768 | 94.10% | 0.78 |
| AG8-Rh89/T | 56,054,626 | 48.56 | 11,793 | 31,591 | 3496.17 | 318,132 | 93.50% | 0.53 |

22

| Table S4. Taxonomic placement of *Rhizoctonia solani* and selected other basidiomycetes genomes used for comparative analysis | | | | | |
|---|---|---|---|---|---|
| **Sub-Division** | **Class** | **Subclass** | **Order** | **Family** | **Species, ENA Accession** |
| Agaricomycotina | Agaricomycetes | | Cantharellales (*Incertae sedis*) | Ceratobasidiaceae | *Rhizoctonia solani* (Teleomorph: *Thanatephorus cucumeris*; soilborne plant pathogens) |
| | | | | Tulasnellaceae | *Tullasnella calospora* (Patch-forming fungus; https://mycocosm.jgi.doe.gov/Tulca1/Tulca1.home .html) |
| | | | Polyporales (*Incertae sedis*) | Fomitopsidaceae | *Postia placenta* (Brown rot fungus; GCA_002117355) |
| | | | | Ganodermaceae | *Ganoderma sinense* (Black Reishi; GCA_002760635) |
| | | | Agaricales | Agaricaeae | *Agaricus bisporus* (Button mushroom; GCA_000300555) |
| | | | | Pleuroptaceae | *Pleurotus ostreatus* (Oyster mushroom; GCA_000697685) |
| | | | | Amanitaceae | *Amanita muscaria* (Fly agaric; GCA_000827485) |
| | | | | Omphalotaceae | *Lentinula edodes* (Shiitake; GCA_002003045) |
| | | | | | *Hypsizygus marmoreus* (Synonym: *H. tessellatus*; White clamshell mushroom; GCA_001605315) |
| | | | | Physalacreaceae | *Armillaria solidipes* (Synonym: *A. ostoyae*; Humongous fungus; Wood or root decays of conifers; GCA_002307675) |
| | Tremellomycetes | | Tremellales | Tremellaceae | *Cryptococcus neoformans* (Teleomorph: *Filobasidiella neoformans*; Animal pathogen; GCA_002215885) |
| | Wallemiomycetes (*incertae sedis*) | | | | *Wallemia ichthyophaga (Halophilic fungus;* GCA_000400465*)* |
| Pucciniomycotina | Pucciniomycetes | | Pucciniales | Pucciniaceae | *Puccinia graminis* (Stem rust; GCA_000149925.1) |
| | | | | Melampsoraceae | *Melampsora larici-populina* (Poplar leaf rust; GCA_000204055.1) |
| Ustilaginomycotina | Exobasidiomycetes | Exobasidi omycetida e | Tilletiales | Tilletiaceae | *Tilletia caries* (Common bunt; GCA_001645005) |
| | Ustilaginomycetes | | Ustilaginales | Ustilaginaceae | *Ustilago maydis* (Corn smut; GCA_000328475.2) |

#### References

1. Amaradasa,B.S., Lakshman,D., Horvath,B.J. and Amundsen,K.L. (2014) Development of SCAR markers and UP-PCR cross-hybridization method for specific detection of four major subgroups of Rhizoctonia from infected turfgrasses. *Mycologia*, 10.3852/13-006.
2. Lakshman,D.K. and Tavantzis,S.M. (1994) Spontaneous appearance of genetically distinct double-stranded RNA elements in Rhizoctonia solani. *Phytopathology*, 10.1094/Phyto-84-633.
3. Cubeta,M.A., Thomas,E., Dean,R.A., Jabaji,S., Neate,S.M., Tavantzis,S., Toda,T., Vilgalys,R., Bharathan,N., Fedorova-Abrams,N., *et al.* (2014) Draft genome sequence of the plant-pathogenic soil fungus Rhizoctonia solani anastomosis group 3 strain Rhs1AP. *Genome Announc.*, 10.1128/genomeA.01072-14.
4. Johnk,J.S., Jones,R.K., Shew,H.D. and Carling,D.E. (1993) Characterization of populations of Rhizoctonia solani AG-3 from potato and tobacco. *Phytopathology*, **83**, 854–858.
5. Papavizas, G. C. and Ayers,W.A. (1965) Virulence, host range, and pectolytic enzymes of single-basidiospore isolates of Rhizoctonia praticola and Rhizoctonia solani. *Phytopathology*, **55**, 111–116.
6. Lewis,J.A. and Papavizas,G.C. (1987) Reduction of inoculum of Rhizoctonia solani in soil by germlings of Trichoderma hamatum. *Soil Biol. Biochem.*, 10.1016/0038-0717(87)90081-2.
7. Lakshman,D.K., Roberts,D.P., Garrett,W.M., Natarajan,S.S., Darwish,O., Alkharouf,N., Pain,A., Khan,F., Jambhulkar,P.P. and Mitra,A. (2016) Proteomic Investigation of Rhizoctonia solani AG 4 Identifies Secretome and Mycelial Proteins with Roles in Plant Cell Wall Degradation and Virulence. *J. Agric. Food Chem.*, 10.1021/acs.jafc.5b05735.

#### Supplementary figures

Figure S1

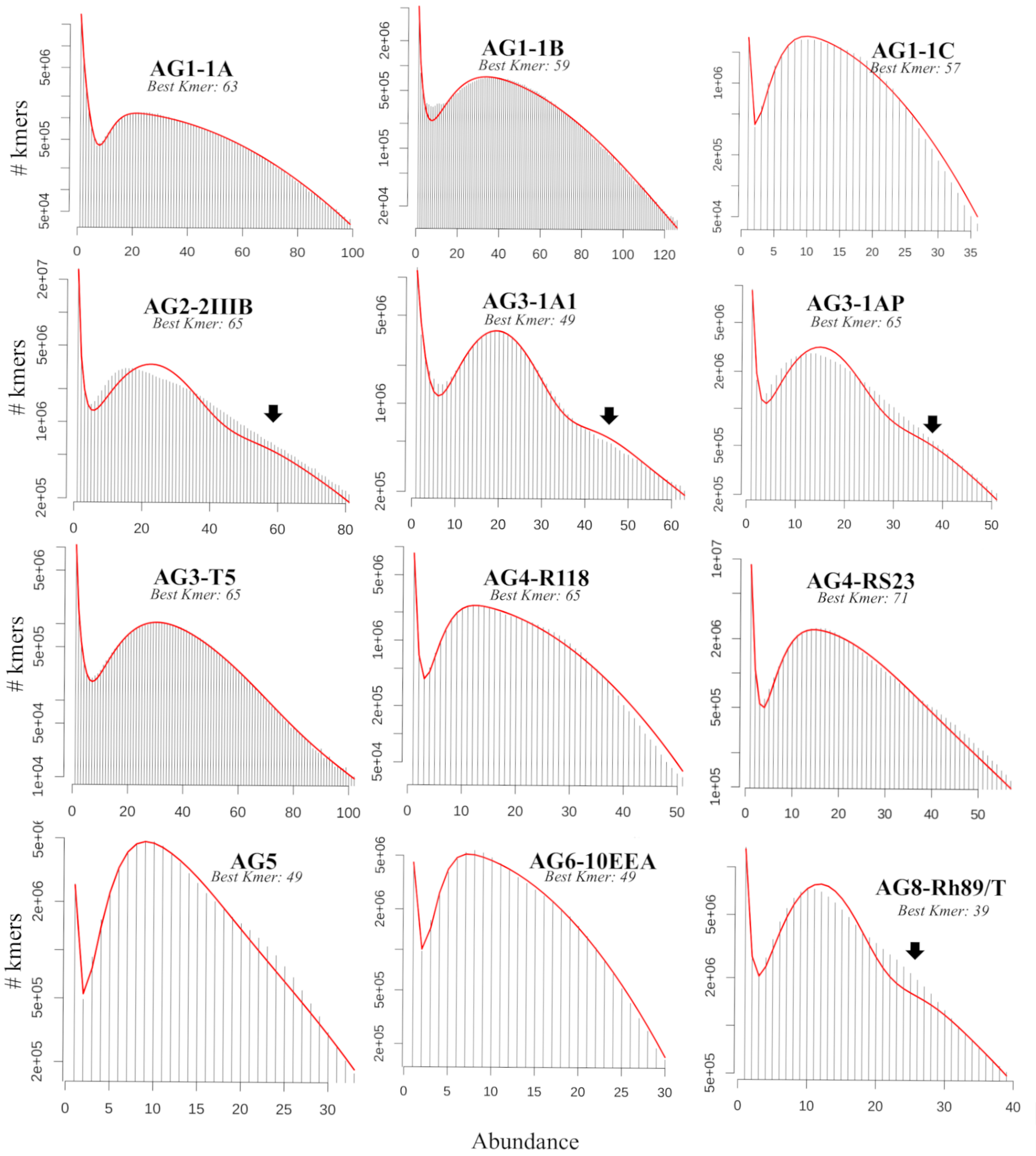

**Figure S1**. *k*-mer distribution. Histogram with the number of k-mers having a given count for the best *k*-mer length, plotted using –histo module of jellyfish program. The x-axis represent number of *k*-mers observed with a given *k*-mer count (y-axis). Arrow marks the shoulder peaks that represent sequence-level heterogeneity or multi-nucleate features in the genome sequencing data.

Figure S2

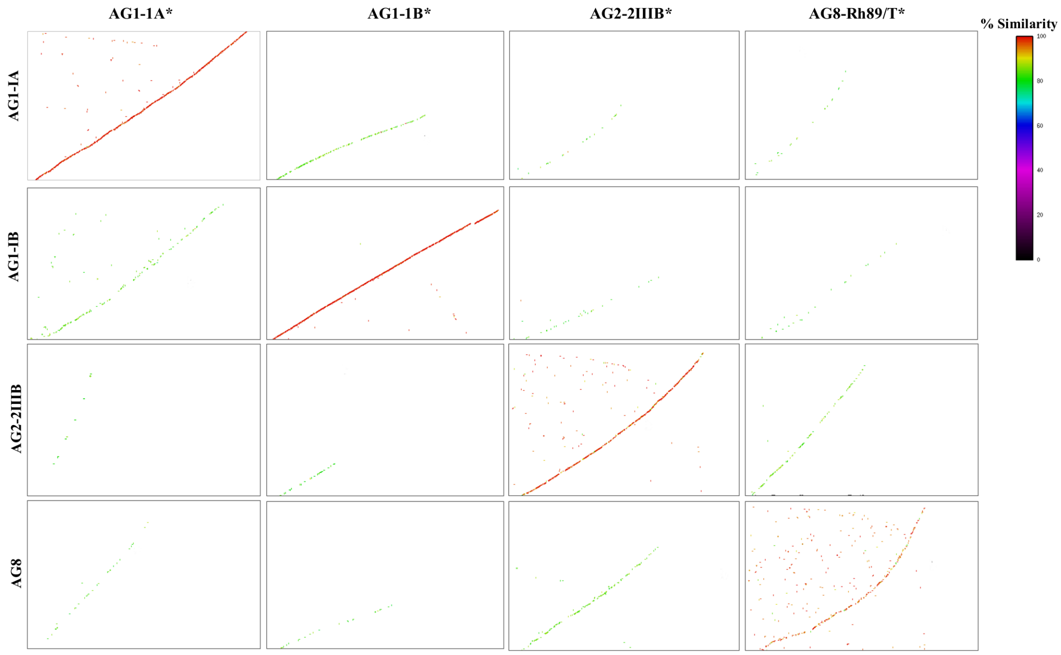

**Figure S2**: Mummer plots of aligned genomic scaffolds of *R. solani* AGs between previously published genomes and de novo genome assemblies (marked with *) produced in this study. The red dots represent high sequence similarity between aligned genomic scaffolds. The scaffolds were aligned using nucmer program and plotted with mummer plot with –filter and –layout arguments.

Figure S3

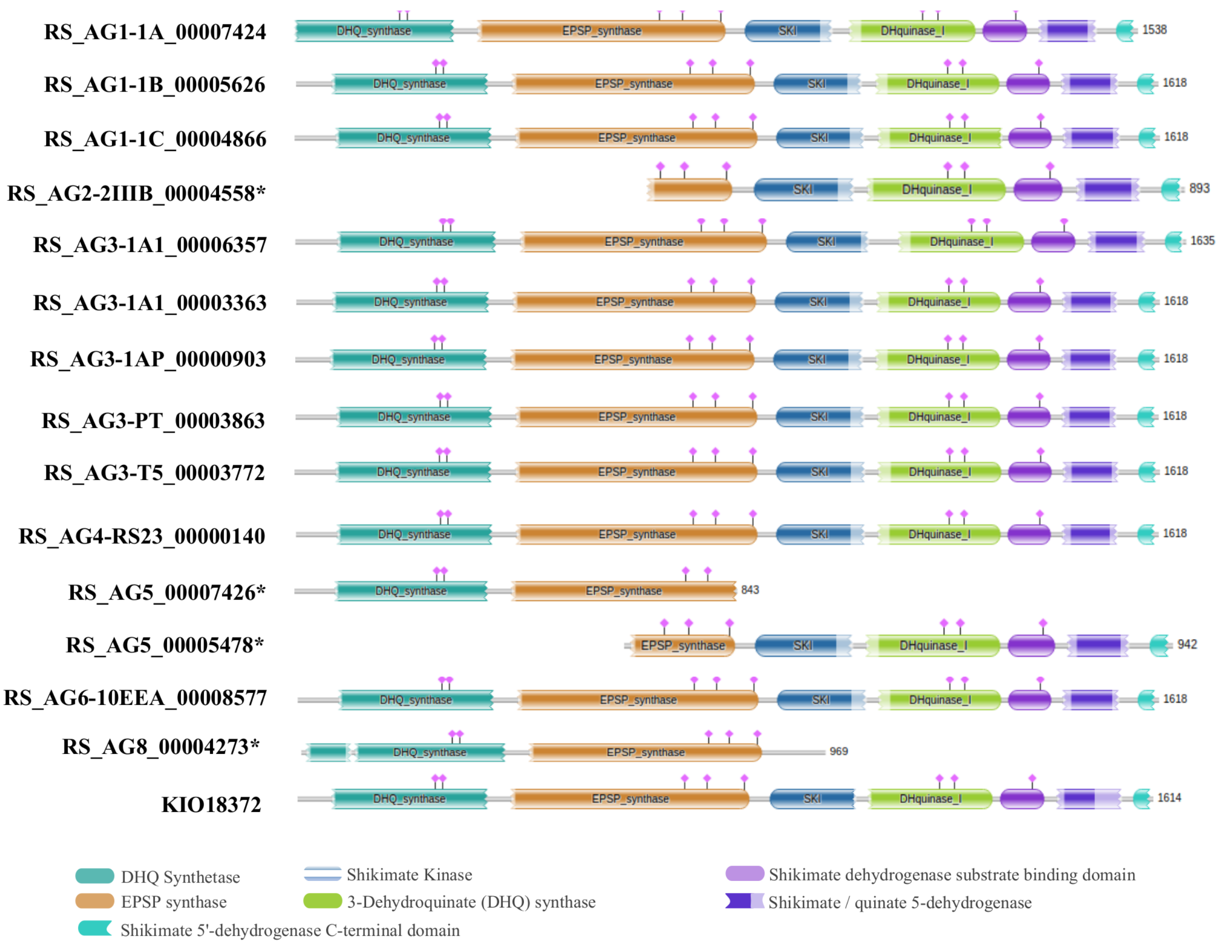

**Figure S3**: A schematic representation of the AROM sequences. The multi-functional AROM sequences predicted in each of the *R. solani* and *T. calospora* genomes are shown. Isolates with partial AROM sequences are marked with *. The domains are predicted and plotted using HMMER webserver (see methods).

Figure S4

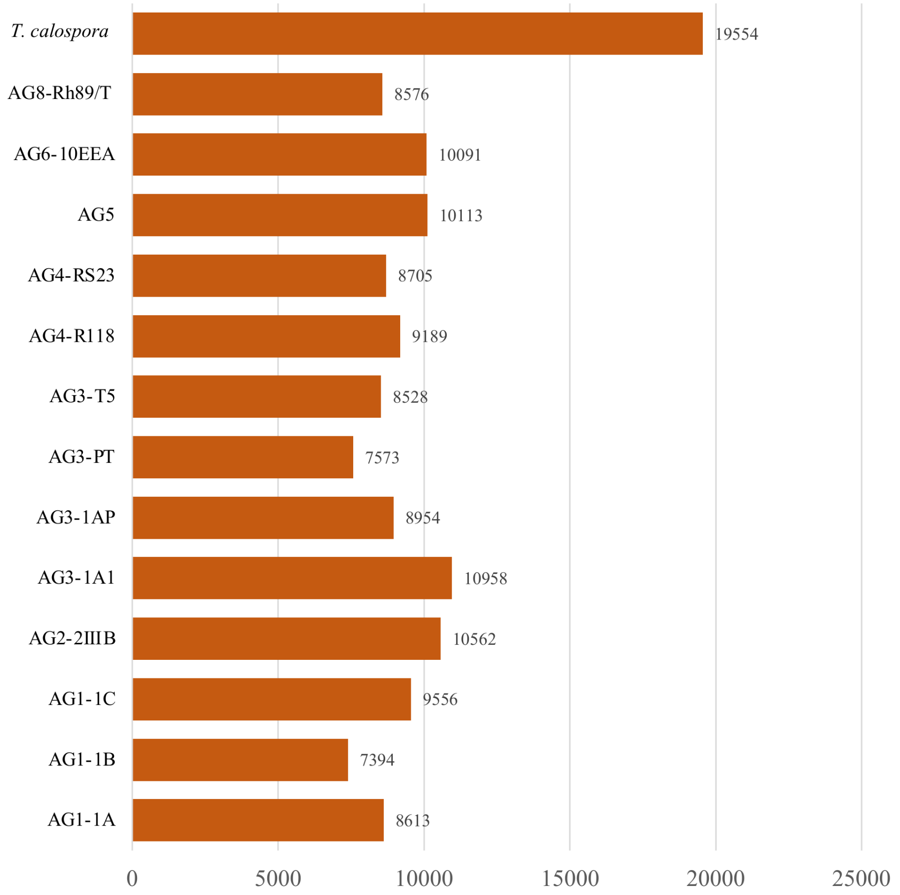

**Figure S4**. The number of protein coding transcripts/proteins predicted in each of the given isolate used in this study.

Figure S5

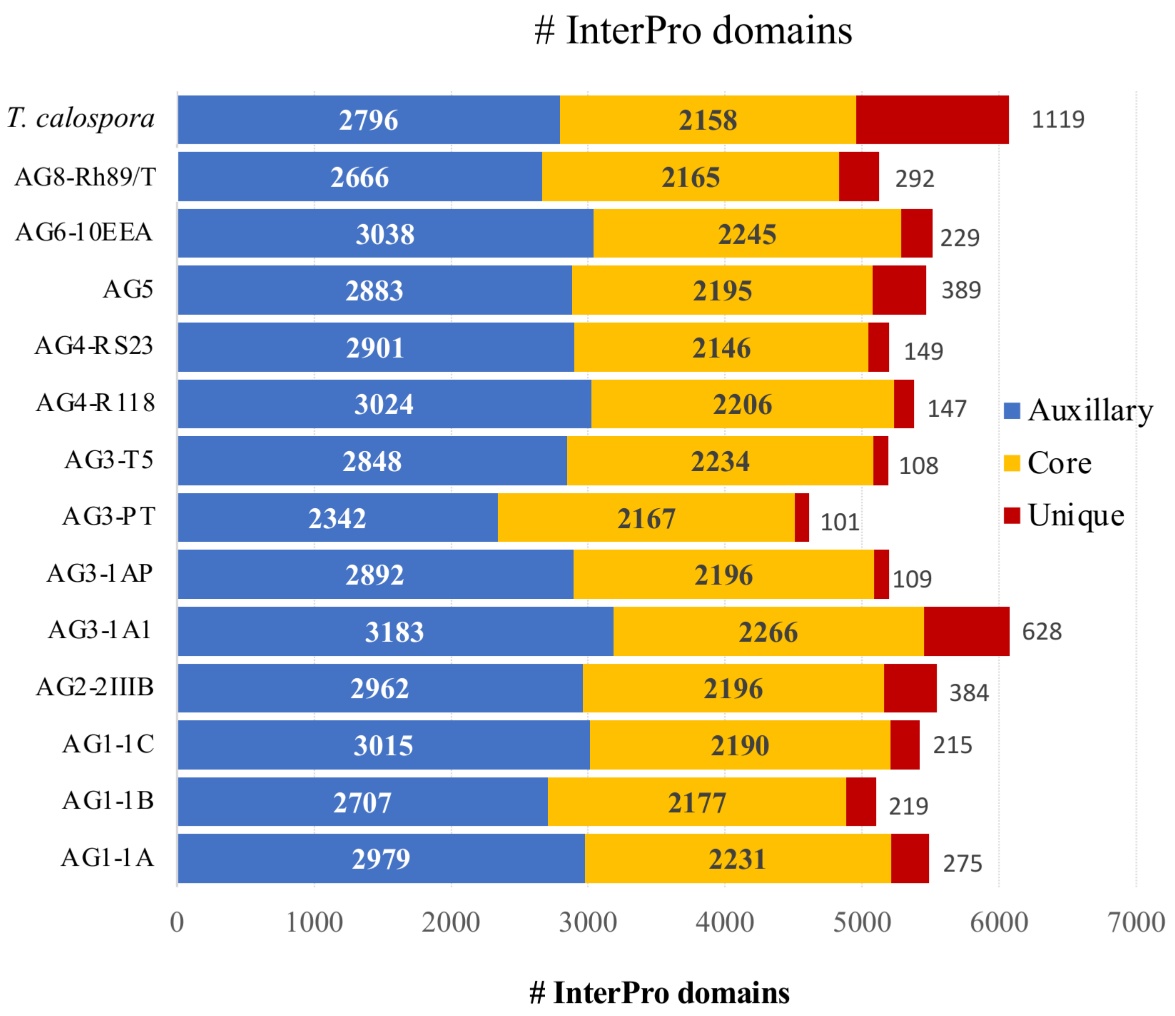

**Figure S5**: **Interpro domain families**. In each isolate, for each type of protein (core, unique and auxiliary), interpro domain families were enumerated. The stacked bar plot shows the number of interpro families enriched with the different protein types in each fungal isolate.

Figure S6

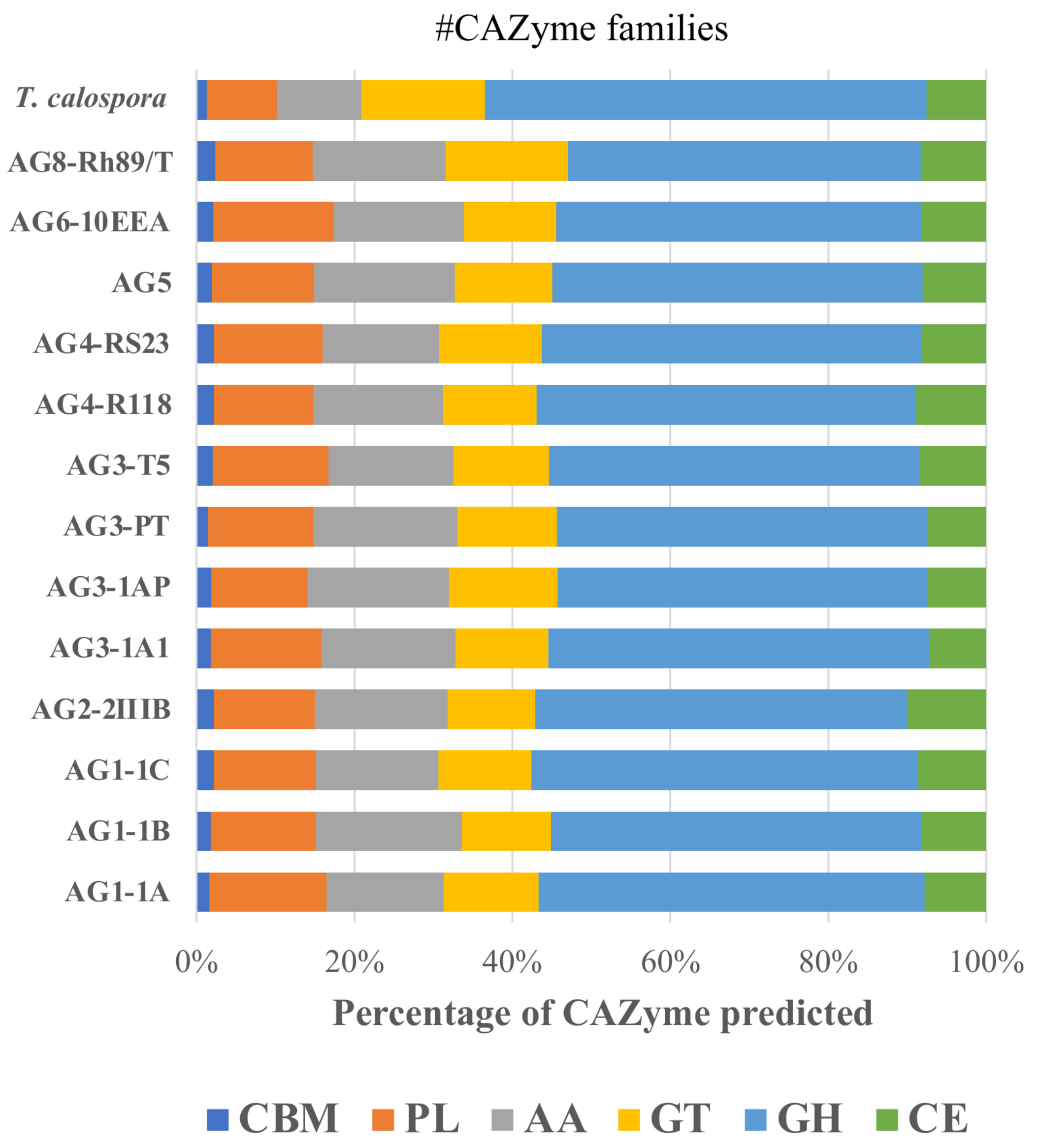

**Figure S6**: CAZyme families. The proportion of proteins, belonging to six different carbohydrate metabolizing families of proteins, predicted in each of the given fungal isolates.

### Figure S7

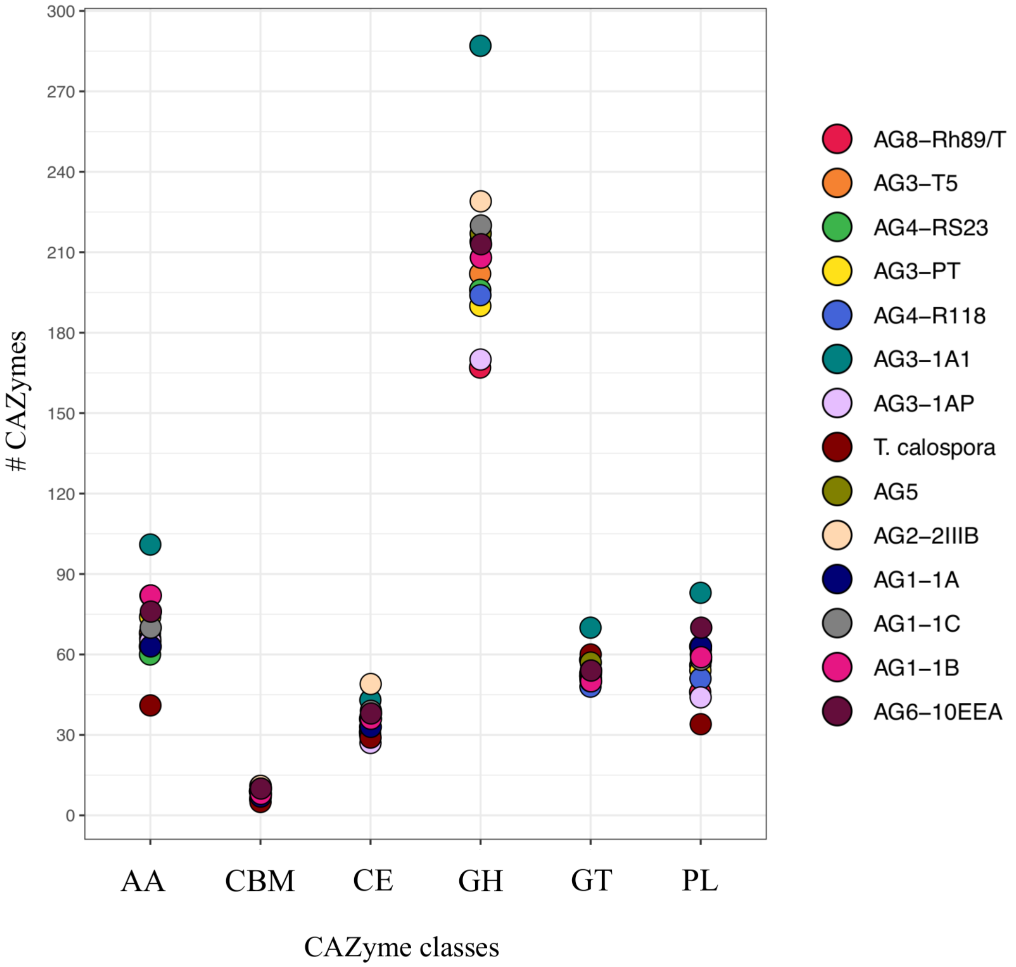

**Figure S7. CAZyme**. The number of CAZymes belonging to different classes, predicted in *R. solani* isolates *and T calospora.*

Figure S8

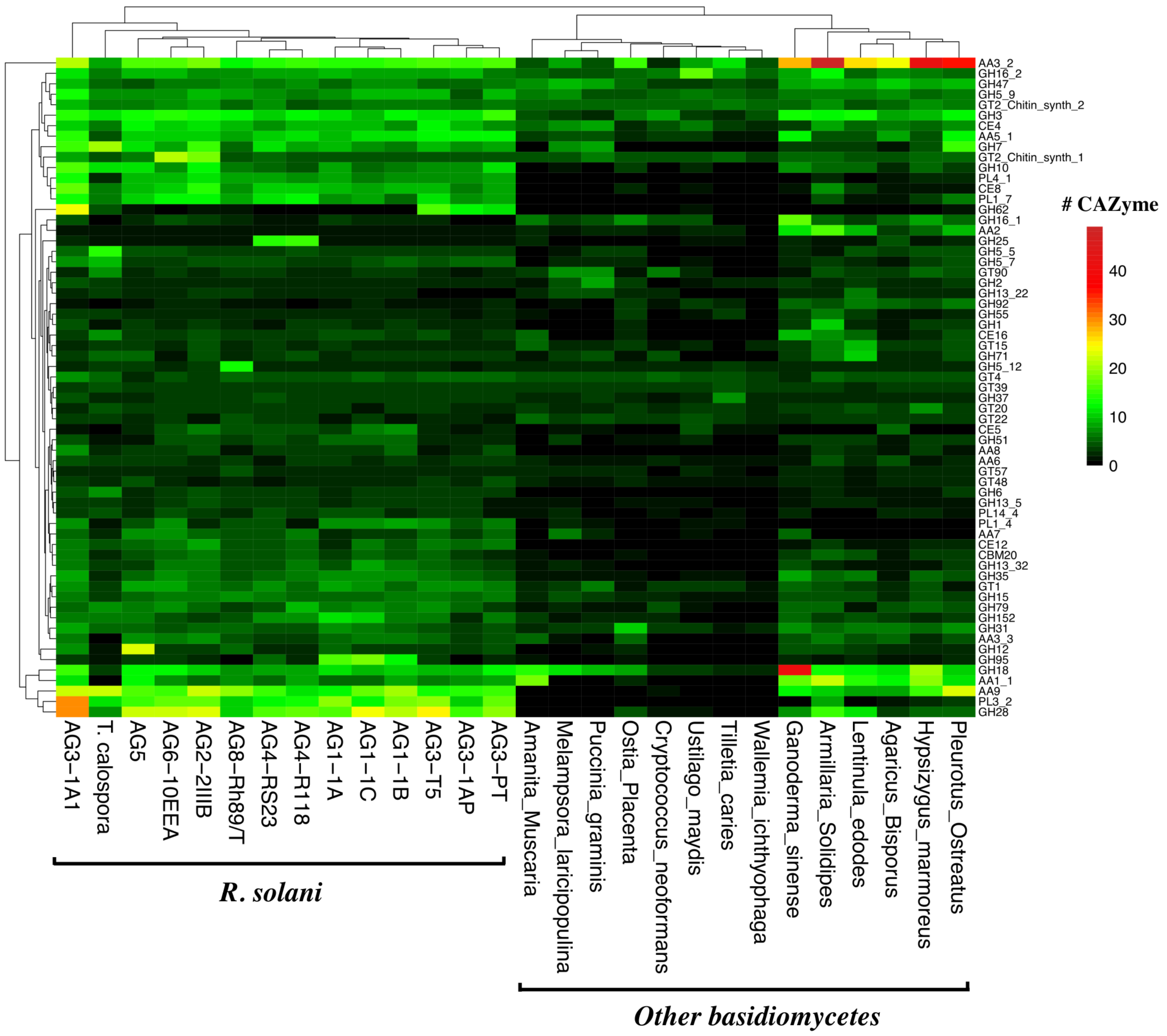

**Figure S8**. **CAZyme**. Heatmap showing the topmost CAZyme family predicted in the proteomes of all the *R solani* AGs, *T calospora* and other basidiomycetes. Each row represents one CAZy family of proteins, and color is proportional to the number of protein member shared in a given family from the given species (black: no member protein; red: large number of members). The phylogenetic analysis enumerates the distance between different fungal isolates based on proteins shared by them across all CAZy families. For simplicity only the CAZyme families enriched in more than 50 proteins across all proteomes were shown.
